# Antisense non-coding transcription represses the *PHO5* model gene at the level of promoter chromatin structure

**DOI:** 10.1101/2022.02.21.481265

**Authors:** Ana Novačić, Dario Menéndez, Jurica Ljubas, Slobodan Barbarić, Françoise Stutz, Julien Soudet, Igor Stuparević

## Abstract

Pervasive transcription of eukaryotic genomes generates non-coding transcripts with regulatory potential. We examined the effects of non-coding antisense transcription on the regulation of expression of the yeast *PHO5* gene, a paradigmatic case for gene regulation through promoter chromatin remodeling. A negative role for antisense transcription at the *PHO5* gene locus was demonstrated by leveraging the level of overlapping antisense transcription through specific mutant backgrounds, expression from a strong promoter *in cis*, and use of the CRISPRi system. Furthermore, we showed that enhanced elongation of *PHO5* antisense leads to a more repressive chromatin conformation at the *PHO5* gene promoter, which is more slowly remodeled upon gene induction. The negative effect of antisense transcription on *PHO5* gene transcription is mitigated upon inactivation of the histone deacetylase Rpd3, showing that *PHO5* antisense RNA acts *via* histone deacetylation. This regulatory pathway leads to Rpd3-dependent decreased recruitment of the RSC chromatin remodeling complex to the *PHO5* gene promoter upon induction of antisense transcription. Overall, the data in this work reveal an additional level in the complex regulatory mechanism of *PHO5* gene expression by showing antisense transcription-mediated repression at the level of promoter chromatin structure remodeling.

## Introduction

The canonical view of eukaryotic transcription has evolved from being considered a highly regulated process initiated from specialized genomic regions, such as gene promoters, to a process that permeates the entire genome (1). In addition to gene promoters, transcription often initiates from intergenic and intragenic regions, as well as regulatory regions such as gene enhancers. Most of the transcripts originating from these regions are non-coding RNAs usually rapidly degraded after synthesis, suggesting that the act of transcription has more potential to exert important biological functions compared to the transcripts themselves (2).

In eukaryotic cells, promoter activation occurs in the context of a repressive chromatin structure, *i.e.* the packing of DNA with histone proteins into nucleosomal arrays (3). Since chromatinized DNA is not accessible for interaction with the transcriptional machinery, activators work in concert with chromatin-modifying and - remodelling factors to expose regulatory sites and allow promoter activation. Chromatin modifiers catalyze covalent modifications of histones, such as acetylation, methylation, and phosphorylation, whereas chromatin remodelers use the energy of ATP hydrolysis to slide histones along the DNA or evict them from the DNA (4, 5). In gene-dense genomes such as that of yeast, transcription often initiates at the 3’ end of genes, leading to the production of antisense (AS) non-coding transcripts (2). AS read-through transcription invades the promoter region of the corresponding gene, where it can exert regulatory effects that are usually repressive to transcription of the coding gene (6–9). Genome-wide and single gene studies have shown that promoters invaded by AS transcription read-through have high nucleosome occupancy and narrow nucleosome-depleted regions (NDRs) (10, 11). Our recent genome-wide study showed that induced elongation of non-coding antisense transcription into coding gene promoters results in increased deacetylation of promoter nucleosomes by Rpd3. Histone deacetylation leads to decreased recruitment of the major chromatin remodeler RSC and consequently to NDR closure, which represses transcription (12). However, there are still few examples of *bona fide* effects of specific AS RNAs on transcriptional regulation of their respective genes, such as the yeast *PHO84* gene.

Studies with the *PHO84* gene have been highly instructive in elucidating the mechanisms of transcriptional regulation through AS non-coding RNAs (13–15). These studies converged on a model in which *PHO84* AS transcription is rapidly terminated in wild-type cells by the NNS (Nrd1-Nab3-Sen1) complex and degraded by the activity of the Rrp6-containing nuclear RNA exosome. Inactivation of any of these crucial factors, such as in *rrp6Δ* mutant cells, leads to transcriptional read-through of *PHO84* AS transcripts, allowing recruitment of histone deacetylases (HDACs) Hda1 or Rpd3 to the *PHO84* promoter. Histone deacetylation is thought to lock the chromatin structure of the promoter in a repressed conformation, thereby negatively regulating transcription of the sense transcript, *i.e. PHO84* mRNA. This mechanism was subsequently explored genome-wide in yeast, which revealed a group of genes that accumulate AS RNAs in the absence of Rrp6 and are silenced in an HDAC-dependent manner (14). Genes of this class are characterized by AS transcripts that span the entire gene length, extend beyond the TSS and are enriched for so-called ‘closed’ promoters. These promoters are typical of inducible or stress-activated genes, and are characterized by precisely positioned nucleosomes whose remodeling is a prerequisite for transcriptional activation (16, 17). A paradigmatic closed promoter that also belongs to this gene class is that of the *PHO5* gene, which is a member of the same (PHO) regulon as *PHO84* (18).

The *PHO5* gene encodes the secreted non-specific acid phosphatase which is located in the periplasmic space and has a role in phosphate metabolism. Accordingly, expression of the *PHO5* gene is regulated in response to intracellular phosphate concentration through the PHO signalling pathway, so that it is repressed when the intracellular concentration is abundant and induced under phosphate starvation conditions (18). This regulation is primarily achieved through phosphorylation of the specific activator Pho4. Under a high phosphate concentration Pho4 undergoes phosphorylation by the cyclin-dependent-kinase Pho80-Pho85, preventing its accumulation in the nucleus and transcriptional activation of the *PHO5* gene. In low phosphate, Pho4 is imported into the nucleus and activates transcription. From the early days of chromatin research in the 1980s until now, the *PHO5* gene promoter has been and continues to be instrumental in the discovery of numerous fundamental principles and mechanisms of chromatin structure remodeling (reviewed in (18)). In the repressed state, the *PHO5* promoter features five precisely positioned nucleosomes, which upon induction are remodelled into a broad nucleosome-depleted region of ∼600 bp. This massive chromatin transition requires the concerted action of a large network of chromatin-modifying and -remodeling complexes as well as histone chaperones. The repressive chromatin conformation is maintained by H3K4 methylation catalyzed by Set1, a mark that recruits the histone deacetylase Rpd3 to the *PHO5* promoter (19, 20). Another histone deacetylase, Hda1, plays a minor role in this process (21). When the intracellular phosphate concentration is limited, signal transduction *via* the PHO signaling pathway leads to the accumulation of the unphosphorylated transcriptional activator Pho4 in the nucleus (18, 22). The first step in transcriptional activation of the *PHO5* gene is the binding of Pho4 to the UASp1 (Upstream activating sequence phosphate 1) site, which is located in the short nucleosome-depleted region between nucleosomes -2 and -3 of the *PHO5* gene promoter. Pho4 recruits histone acetyltransferases, such as the Gcn5-containing SAGA complex, which establish a hyperacetylated promoter configuration (23, 24). Acetylated histones are read by chromatin-remodeling complexes containing bromodomains (25, 26). Alternatively, these remodelers can be recruited to the *PHO5* promoter by direct interaction with Pho4 (27). Five remodelers (SWI/SNF, RSC, INO80, Isw1, Chd1) from all four yeast remodeler families cooperate to catalyze the chromatin opening at the *PHO5* promoter (28, 29), with the most abundant remodeler, RSC, providing the crucial share of the remodeling activity required for this transition (29). Histone eviction allows Pho4 to bind to the UASp2 site otherwise covered by nucleosome -2, which is ultimatively required for full transcriptional activation (30–32).

Another level of *PHO5* promoter regulatory complexity was revealed upon mapping of the *PHO5* AS transcript, CUT025 (33, 34). This transcript initiates from the 3’ region of the *PHO5* ORF and extends through its promoter region, spanning ∼2.4 kb in size. It is produced only in cells growing under repressive (phosphate-rich) conditions and is more abundant in *rrp6Δ* mutant compared to wild-type cells, indicating its degradation by the nuclear RNA exosome (33). AS transcription is generally associated with a repressive effect on transcription of the corresponding genes, and the *PHO5* gene is among the rare examples for which AS transcription is proposed to have a positive effect (33). In this work, we examined the effect of non-coding AS transcription on *PHO5* gene expression by enhancing or impairing elongation of the *PHO5* AS transcript. In both cases, our results argue in favour of antisense transcription having a negative effect on *PHO5* gene expression. Moreover, we provide evidence that this negative effect occurs through a chromatin-remodeling based mechanism mediated by AS transcription which decreases the accessibility of the chromatin structure at the *PHO5* gene promoter.

### Materials and methods

Yeast strains and primer sequences used in this study are listed in Supplementary Tables S1 and S2, respectively.

### Strains, media, plasmids and strain construction

Yeast *Saccharomyces cerevisiae* strains used in this study are listed in Supplementary Table S1. All strains were grown at 30°C. For repressive conditions (high phosphate, +P_i_), yeast strains were grown in YNB medium supplemented with 1 g/l KH_2_PO_4_ (YNBP) with or without lack of amino acids for plasmid selection. For gene induction by phosphate starvation (-P_i_), cells were washed in water and resuspended in the phosphate-free synthetic medium with or without lack of amino acids for plasmid selection (28, 29, 35). Anchor-away of Nrd1-AA and Sth1-AA was induced by adding 1 µg/ml of rapamycin (Sigma) to the medium. The *RRP6* gene was deleted using a disruption cassette generated by PCR with the primer pairs RRP6-Kan1 and RRP6-Kan2 or RRP6hph_fwd and RRP6hph_rev and the BMA41 *rrp6::KanMX4* genomic DNA or the *hph*-carrying pYM16 plasmid from (36) as template, respectively. The *GCN5* gene was deleted using a disruption cassette generated by PCR with the primer pair gcn5HIS_fwd and gcn5HIS_rev and the *SpHIS5*-carrying pKT101 plasmid from (37) as template. Transformants were selected on G-418 (0.2 mg/ml, Sigma), Hygromycin B (0.3 mg/ml, Sigma) or -His plates, depending on the marker, and gene deletion was confirmed by PCR. The BMA41 *TEF1-PHO5 AS* strain was constructed by transformation with a cassette generated by PCR with primers TEF1PHO5AS_fwd and TEF1PHO5AS_rev and the pYM-N18 plasmid from (36) as template. Transformants were selected on G-418 plates, and correct insertion of the cassette was confirmed by PCR. The pP5Z reporter plasmid is centromeric vector that carries a *PHO5* promoter-*lacZ* gene fusion and is described in (38). The pCEN-RRP6 plasmid was previously constructed by Gateway cloning from the pAG416GPD backbone (39). Plasmid pTDH3-dCas9 (pFS3891) (40) was obtained from Addgene (Plasmid #46920). Plasmid pFS3892, which contains the guide RNA scaffold, was generated by one-step isothermal Gibson assembly reaction (New England BioLabs) using two fragments, one obtained by PCR on pRPR1_gRNA_handle_RPR1t (Addgene Plasmid #49014) using OFS_2869 and OFS_2870 oligonucleotides, the other by PCR on YCpLac33 using OFS_2871 and OFS_2872 oligonucleotides. Plasmid *PHO5 AS gDNA-URA3* was then obtained by Gibson assembly reaction (NEB) using OFS_2886 and OFS_2887 to amplify pFS3892 backbone and OFS_2888 and OFS_3095 for gDNA cloning. To test the putative *in trans* activity of the *PHO5* AS RNA (Figure 4C), strains were designed as following. First, the *PHO5* ORF was replaced by the *URA3* marker in either a *MATA* FSY6857 or a *MATα* FSY5439 strain (see Table S1). This was performed by amplification of the *URA3* marker from the pUG72 plasmid with OFS5084 and OFS5085 primers and the resulting amplicon was transformed in FSY6857 and FSY5439 strains following selection in a -Ura medium. The *MATA* and *MATα* strains deleted for *PHO5* were named FSY9286 and FSY9287. We then amplified the wild-type *PHO5* gene with either the OFS5086 and OFS5087 primer pair or OFS5088 and OFS5089 primer pair in order to insert a terminator for the *PHO5* mRNA (sense) or the AS transcript, respectively. The PCR products targeting either the sense or the antisense transcription were transformed in the FSY9286 and FSY9287 and counter-selected on a 5-FOA medium. The strains targeting the *PHO5* mRNA or AS RNA were named FSY9288 and FSY9291. Finally, the 3 different diploids (Figure 4C: AS in *cis*, AS blocked and AS in *trans*) were generated after crossing FSY6857 with FSY9287, FSY9286 with FSY9291 and FSY9288 with FSY9291, respectively, and selection on -His-Trp medium.

### Enzyme activity assays, RNA isolation, Nothern blot and RT-qPCR

Acid phosphatase and beta-galactosidase activity assays were done with intact yeast cells, exactly as described in (29). Total RNA was extracted by the hot phenol method (41), treated with RNAse-free DNAse I (New England Biolabs) and purified by phenol/chloroform extraction. Strand-specific reverse transcription was performed using 1 µg of RNA and strand-specific oligonucleotides (0.1 µM each) with the ProtoScript First Strand cDNA Synthesis Kit (New England Biolabs) supplemented with actinomycin D (Sigma) to final concentration 5 μg/ml to ensure strand specificity. cDNAs were amplified in Roche LightCycler 480 with the Maxima SYBR Green qPCR Master Mix detection kit (Thermo Scientific). The qPCR datasets were analysed using the ΔΔCt method, and the results were normalized to either *PMA1, ACT1* or *SCR1* RNAs amplification, which were used as internal controls. For Figure 4C, OFS2522 and OFS2523 were used to measure *PHO5* mRNA and AS RNA levels. Amplifications were done in duplicate for each sample, and three independent RNA extractions were analysed. For the Northern blot, total RNA (10 μg for each sample) was run on a 1% denaturing formaldehyde agarose gel and transferred to nylon membranes (Amersham Hybondtm-N+). Membranes were crosslinked and incubated overnight at 60°C with 100μg/ml boiled salmon sperm DNA in 50% formamide, 5x standard saline citrate (SSC), 20% dextran sulfate sodium, 1% sodium dodecyl sulfate (SDS). Subsequently, membrane wered hybridized with 32P-labeled SP6/T7 riboprobes in 50% formamide, 7% SDS, 0.2 M NaCl, 80 mM sodium phosphate (pH 7.4), and 100 μg/ml boiled salmon sperm DNA for 6h. All blots were washed with 2X SSC and 0.1% SDS for 5 minutes at60°C and then with 0.5X SSC and 0.1% SDS for 45 minutes at 60°C. Riboprobes were obtained by SP6/T7 in vitro transcription of gene-specific PCR fragments containing an SP6/T7 promoter. Quantifications were performed with a Phosphor Imager machine.

### Chromatin analysis

For anti-histone H3 chromatin immunoprecipitation (ChIP), forty millilitres of cells were fixed with 1% formaldehyde for 20 min. After quenching with 400 mM glycine to stop the reaction, the cells were washed and lysed with glass beads to isolate chromatin. Sonication of cell lysates was performed with a Vibra-Cell sonicator in 1.2 mL of FA150 buffer (50 mM Hepes pH 7.5, 150 mM NaCl, 1 mM EDTA, 1% Triton X-100, 0.1% Sodium deoxycholate and 0.1% SDS) to reduce average fragment size to approximately 500 base pairs. The samples were centrifuged at 2500 g and the supernatant recovered. Chromatin fractions of 400 μl were taken for each immunoprecipitation reaction and incubated with 4 μl of anti-histone H3 antibodies (ab1791, Abcam) at 4°C overnight. After incubation, 40 μl of protein G PLUS-agarose beads (sc-2002, Santa Cruz Biotechnology) were added and incubated at 4°C for 2 h.

The beads were washed extensively by successive washing steps: 3 times with FA150 lysis buffer, 3 times with FA500 lysis buffer (similar to FA150 but with 500 mM NaCl), 1 time with washing buffer 1 (10 mM Tris-HCl pH 8, 250 mM LiCl, 1 mM EDTA, 0.5% NP-40, 0.5% Sodium deoxycholate) and 1 time with washing buffer 2 (10 mM Tris-HCl pH 8, 1 mM EDTA, 1% SDS). Chromatin was eluted at 80°C in elution buffer (50 mM Tris-HCl pH 8, 10 mM EDTA, 1% SDS) during 20 minutes. Samples (regardless of Input or IP) were reverse cross-linked at 65°C overnight. Eluted supernatants (output) and the input controls were hydrolysed with Pronase (0.8 mg/ml final concentration, Sigma) at 42°C for 2 h, followed by incubation at 65°C for 7 h to reverse cross-linked DNA complexes. DNA was extracted using the Macherey Nagel Nucleospin Gel & PCR Cleanup Kit. The immunoprecipitated DNAs (output) were quantified by qPCR in Roche LightCycler 480 with the Maxima SYBR Green qPCR Master Mix detection kit (Thermo Scientific). Amplifications were done in triplicate for each sample. Immunoprecipitated samples (output) were normalized to input and to a *PHO5*- adjacent control region which does not show chromatin signatures similar to the *PHO5* gene promoter, as described in (31). Chromatin analysis of yeast nuclei by restriction nuclease accessibility assay was done as described previously (29, 35, 42). 120 U of the ClaI restriction enzyme (New England Biolabs) was used for chromatin analysis of nuclei and 40 U of HaeIII (New England Biolabs) was used for secondary cleavage. Probe for hybridization was as described previously (29, 35, 43). Quantification of the percentage of cleaved DNA was done by PhosphorImager analysis (Fuji FLA3000). ChIP of dCas9 was essentially perfomed as in (12) without addition a *S. pombe* spike-in. An anti-Cas9 antibody (Diagenode #C15310258) was used for the immunoprecipitation step.

### Downloaded data sets

For RNA-seq and RNAPII PAR-CLIP, data were retrieved from (44) (GEO: GSE175991) and from (45) (GEO: GSE56435). Data of MNase-seq, ATAC-seq and Sth1 ChEC-seq were reanalyzed from (12) (GEO: GSE130946).

## Results

### AS transcription is involved in regulation of *PHO5* gene expression

The product of antisense transcription at the *PHO5* model gene locus, CUT025 (hereafter referred to as *PHO5* AS), is initiated at the 3’ end of the gene ORF in the antisense direction and extends through the *PHO5* promoter region (Fig. 1A). The 3’-5’ exoribonuclease Rrp6, which is the catalytic subunit of the nuclear RNA exosome complex, degrades this transcript in wild-type (wt) cells, consistent with the increased level of this transcript in *rrp6Δ* mutant cells (Fig. S1A). We confirmed the increased level of the *PHO5 AS* transcript at the *PHO5* promoter region in *rrp6Δ* compared to wt cells by strand-specific reverse-transcription quantitative PCR (RT-qPCR) upon shifting the cells from repressive (phosphate-rich, +P_i_; YNB with additional 1 g/l KH_2_PO_4_) to inducing (no phosphate, -P_i_) conditions. *PHO5* AS accumulation in *rrp6Δ* was most pronounced under repressive conditions (Fig. 1A, 0 h of induction), consistent with (33). After shifting to inducing conditions, the level of *PHO5* AS gradually decreased in both wild-type and *rrp6Δ* cells, however the increased level in *rrp6Δ* cells was still present at an early time point of gene induction (Fig. 1A). The *PHO5* AS transcript has a much lower steady-state level than the corresponding *PHO5* mRNA transcript, as observed by RNA-seq, which measures steady-state RNA levels, *i.e.*, takes into account both the level of nascent transcription and RNA degradation. However, the RNAPII photoactivatable ribonucleoside-enhanced crosslinking and immunoprecipitation (PAR-CLIP) signal, which measures only nascent transcription, is comparable or even higher for the AS transcript than for the mRNA transcript under same growth conditions (4 mM P_i_), showing that the AS transcript is being produced to a potentially significant level (Fig. 1B). Whole-genome tiling array datasets revealed production of another non-coding transcript at the *PHO5* gene locus, SUT446, transcribed in the sense direction through the *PHO5* promoter region, which appears not to be accumulated in *rrp6Δ* mutant cells and is weakly expressed ((14, 34); Fig. S1A). It was determined by RT-qPCR that the level of SUT446 was not significantly increased in *rrp6Δ* compared to wild-type cells neither in repressive nor inducing conditions (Fig. S1B), arguing against its gene-regulatory function. Overall, these data support a possible regulatory role of the CUT025 AS non-coding transcript, but not the SUT446 promoter non-coding transcript, in regulation of *PHO5* gene transcription.

**Figure 1.**
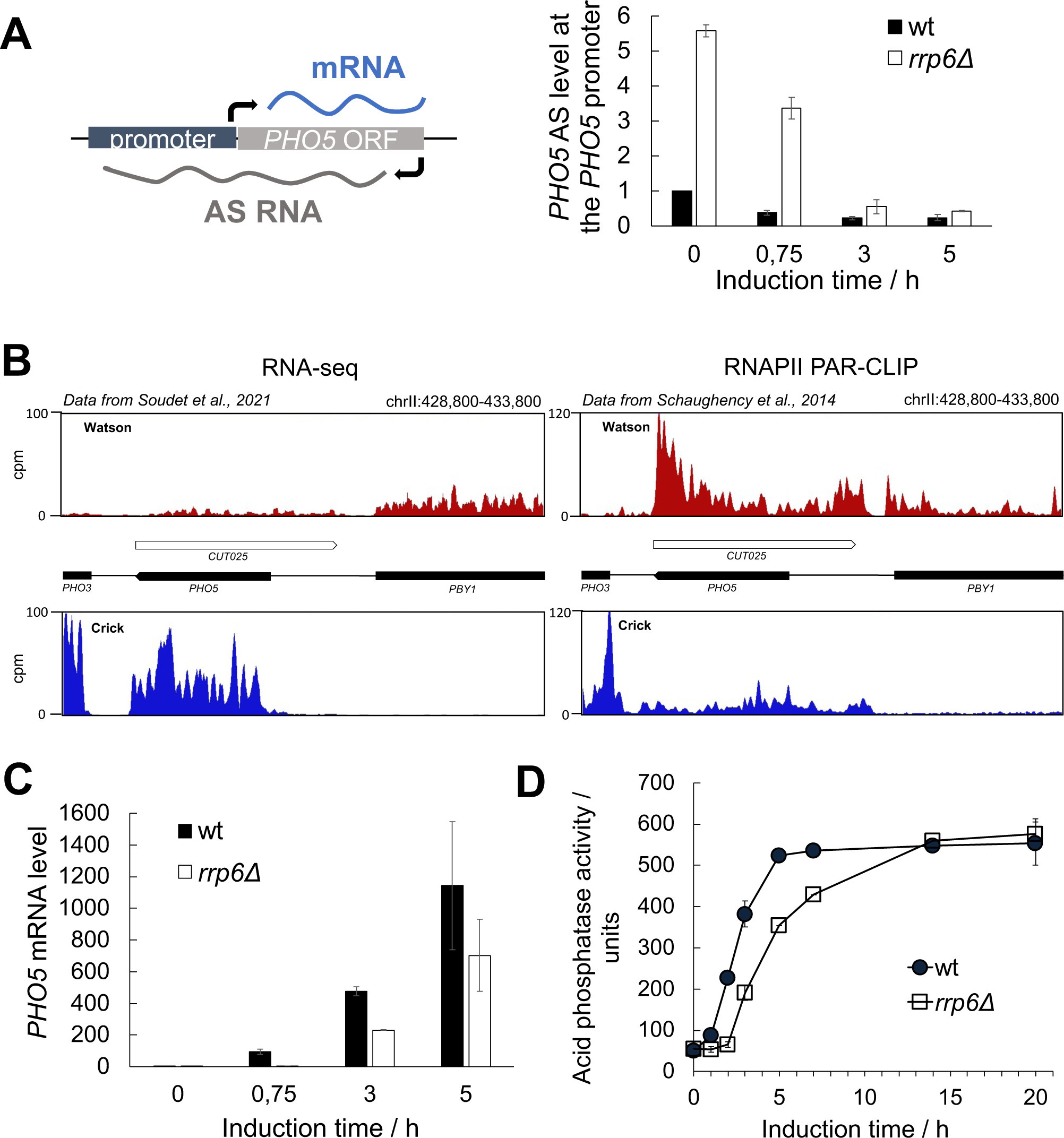
Kinetics of *PHO5* gene expression are inversely correlated with level of the corresponding antisense transcript. **(A)** Scheme showing transcription of an antisense (AS) RNA at the *PHO5* gene locus (left) and its levels at the *PHO5* promoter region in wild-type BMA41 (wt) and corresponding *rrp6Δ* mutant cells upon induction through phosphate starvation, monitored by strand-specific reverse transcription quantitative PCR (RT-qPCR) (right). RT-qPCR values were normalized to *PMA1* RNA and expressed relative to transcript abundance in wild-type cells under repressive conditions (0 h of induction), which was set to 1. **(B)** The left panel shows RNA-seq signal from an Nrd1-AA strain in the absence of rapamycin (wild-type equivalent) at the *PHO5* locus. The right panel represents RNAPII PAR-CLIP signal or nascent transcription signal in the same conditions. Data were retrieved from (44) and (45), respectively. **(C)** Levels of *PHO5* mRNA in wild-type BMA41 (wt) and corresponding *rrp6Δ* mutant cells upon induction through phosphate starvation. RT-qPCR values were normalized to *PMA1* RNA and expressed relative to transcript abundance in wild-type cells at repressive conditions (0 h of induction), which was set to 1. **(D)** Same as (C), but acid phosphatase induction kinetics were monitored by measuring acid phosphatase activity with whole cells. Reported values represent the means and standard deviations of three independent experiments (n = 3).

We further investigated whether the increased level of the *PHO5* AS transcript under repressive conditions and during early gene induction in *rrp6Δ* cells correlates with a change in *PHO5* mRNA level. *PHO5* mRNA was quantified by RT-qPCR upon gene induction and a strong delay in its expression was observed in *rrp6Δ* cells compared to wild-type cells (Fig. 1C). This delay persisted during the first hours of gene induction and corresponded to a delay in expression of the Pho5 acid phosphatase, as determined by measuring its enzymatic activity (Fig. 1D). However, after prolonged induction, the level of acid phosphatase in *rrp6Δ* cells reached that of wild-type cells (Fig. 1D). The observed delay in gene expression was dependent on the catalytic activity of Rrp6, because the catalytically dead *rrp6Y361A* mutant cells also exhibited delayed *PHO5* gene expression, and acid phosphatase activity was brought to wild-type levels when a functional *RRP6* gene was expressed from a centromeric plasmid in *rrp6Δ* cells (Fig. S2A). A similar delay was also measured with *rrp6Δ* cells of two other genetic backgrounds (Figs. S2B and S2C), showing that it is not specific to the W303-derived strain used in these experiments. We also performed a control experiment to test whether the observed kinetic delay in *PHO5* expression in *rrp6Δ* cells is caused by an indirect effect due to compromised signal transduction through the PHO signaling pathway. We made use of a construct in which expression of the *lacZ* reporter gene was driven by the *PHO5* promoter and monitored its expression by measuring beta-galactosidase activity upon induction (no phosphate, - P_i_) in wild-type and *rrp6Δ* cells (Fig. S2D). Expression kinetics of the *PHO5* promoter-*lacZ* construct were similar in wild-type and *rrp6Δ* cells, arguing that PHO signaling is not compromised in *rrp6Δ* cells. This result demonstrates that the kinetic delay in *PHO5* expression observed with the *rrp6Δ* strain (Figs.1C and D) was not an indirect effect caused by compromised induction strength and consequently impaired *PHO5* transcriptional activation. Additionally, this result speaks in favour of a possible regulatory role of the AS transcript originating from the *PHO5* ORF.

### *PHO5* gene expression kinetics are delayed upon induction in mutants related to RNA exosome function

Rrp6 is the nuclear-specific catalytic subunit of the RNA exosome complex. To determine the involvement of other RNA exosome subunits and cofactors in the regulation of *PHO5* gene expression, we examined the kinetics of *PHO5* gene expression using appropriate mutant cells. Deletion mutants for the monomeric cofactors of the nuclear exosome, Rrp47 and Mpp6, also showed delayed acid phosphatase expression kinetics (Fig. 2A). The TRAMP complex is another cofactor of the nuclear RNA exosome and consists of a non-canonical poly(A) polymerase (Trf4 or Trf5), an RNA-binding subunit (Air1 or Air2), and the essential helicase Mtr4 (46, 47). Interestingly, single *air1Δ* and *air2Δ* mutant cells showed no delay, whereas the *air1Δair2Δ* double mutant showed an even greater delay than the *rrp6Δ* mutant (Fig. 2A), consistent with a high degree of redundancy between homologous TRAMP subunits (48). Somewhat surprisingly, acid phosphatase activity measured after overnight induction was increased in some mutants compared with wild-type cells. It is possible that this may reflect specialized cofactor requirements that support the specific conditions of prolonged gene induction and was not pursued further. The mutant for the exonuclease activity of the essential RNA exosome catalytic subunit Dis3 (*dis3Δ + pDis3-exo^-^*) also showed delayed kinetics compared with the corresponding wild-type cells (*dis3Δ + pDis3*) and with the mutant for its endoribonuclease activity (*dis3Δ + pDis3-endo^-^*) (Fig. 2B). These results demonstrate the involvement of the second catalytic subunit of the RNA exosome, Dis3, as well as the nuclear RNA exosome cofactors Rrp47, Mpp6 and the TRAMP complex in the regulation of *PHO5* gene expression.

**Figure 2.**
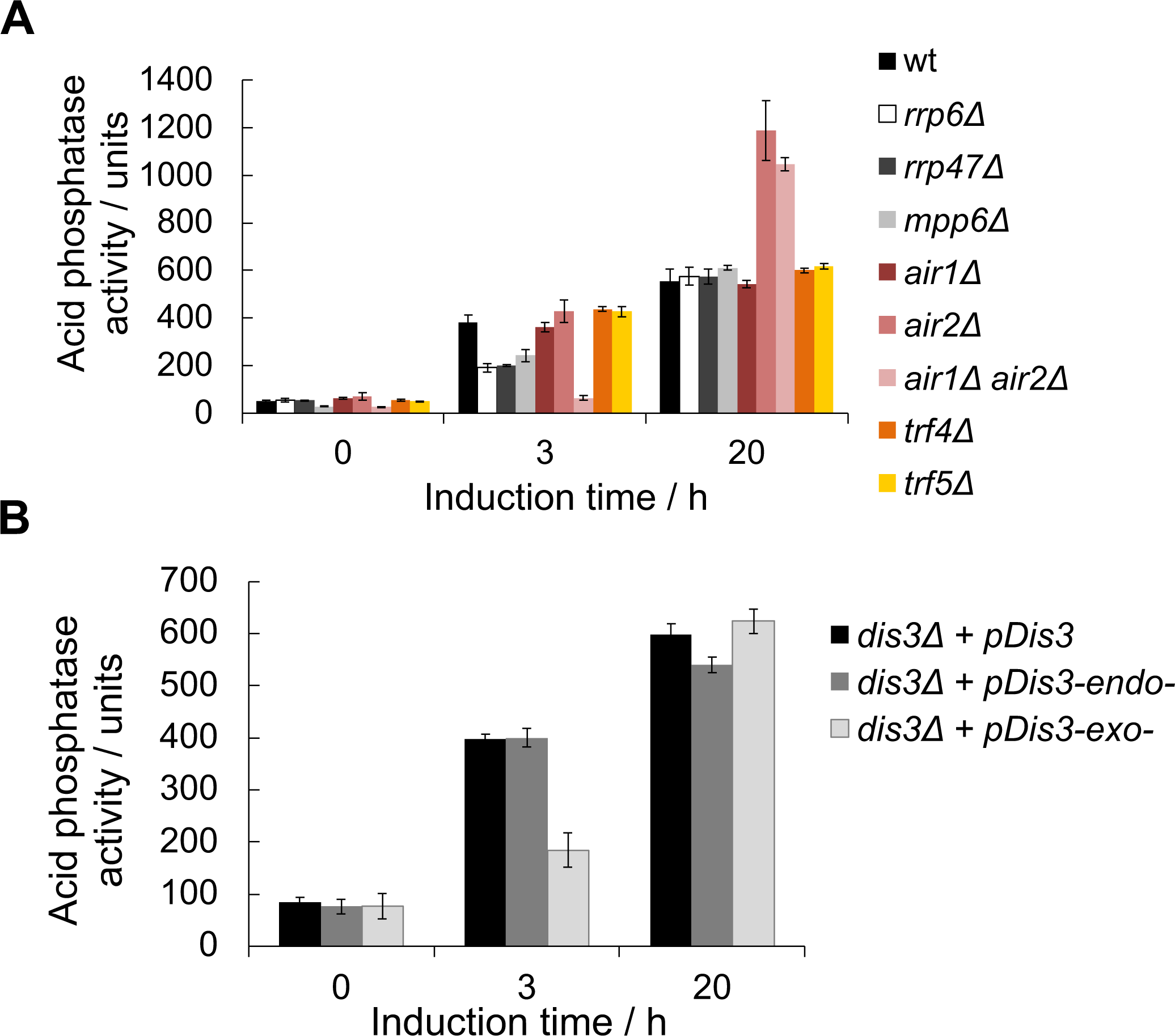
Expression of the *PHO5* gene is negatively affected in RNA exosome mutant cells. **(A)** Acid phosphatase induction kinetics in wild-type BMA41 (wt) and corresponding deletion mutant cells for Rrp6 and RNA exosome cofactors upon induction through phosphate starvation. Reported values represent the means and standard deviations of three independent experiments (n = 3). **(B)** Same as (A), but for W303-derived strains with genomic copy of *DIS3* gene deleted but bearing a centromeric plasmid that carries the wild-type copy of *DIS3* gene (*dis3Δ + pDis3*) or its alleles with abolished endonuclease (*dis3Δ + pDis3-endo^-^*, D171N) or exonuclease (*dis3Δ + pDis3-exo^-^*, D551N) activity.

In *rrp6Δ* and other RNA exosome deletion mutant backgrounds, AS transcription is constitutively induced due to sequestration of the NNS (Nrd1-Nab3-Sen1) termination complex by stabilised non-coding RNAs. The NNS complex cannot be efficiently recycled to sites of transcription, inducing termination defects at non-coding RNA loci and resulting in their increased elongation frequency (49). To rule out possible indirect effects on transcription of the *PHO5* gene due to gene deletion mutant backgrounds in which AS transcription is constitutively elongated, we turned to a system in which AS elongation is inducible. To this end, we used the Anchor Away (AA) system to rapidly deplete Nrd1 protein from the nucleus by rapamycin treatment (50). Since Nrd1 belongs to the NNS surveillance system, its removal is expected to trigger transcriptional read-through of non-coding RNAs (49). Indeed, treatment of Nrd1-AA cells with rapamycin resulted in rapid induction of the *PHO5* AS transcript production, clearly demonstrating that the NNS complex is important for its early termination in wild-type cells. Importantly, even under *PHO5* repressive conditions, induction of *PHO5* AS transcript production through the Nrd1-AA system was accompanied by downregulation of *PHO5* mRNA levels, as shown by Northern blot (Fig. 3A). Furthermore, with the Nrd1-AA system, it was possible to induce elongation of AS transcription by adding rapamycin simultaneously when shifting the cells to *PHO5* inducing conditions (*i.e.* phosphate free medium) (Fig. 3B) or an hour before the shift (Fig. 3C). Consistently, the kinetics of Pho5 expression monitored by measuring acid phosphatase activity showed a kinetic delay which was dependent on the timing of rapamycin addition during cultivation (Figs. 3B and 3C). The results of this experiment demonstrated that the negative correlation between *PHO5* AS and mRNA transcript levels is not an indirect consequence of gene deletion mutant backgrounds, since it is also seen upon induced Nrd1 depletion.

**Figure 3.**
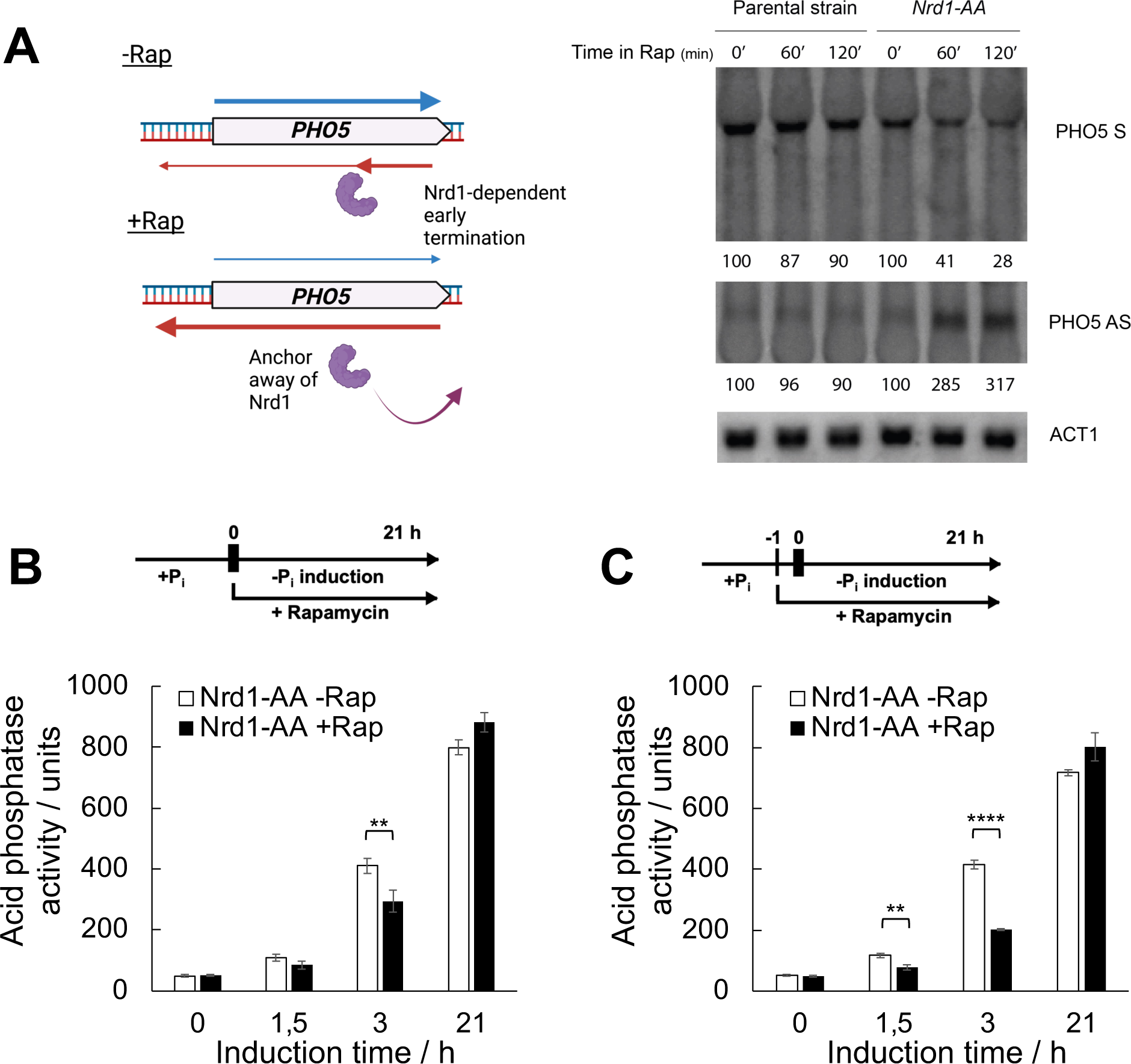
Induction of *PHO5* AS elongation by depletion of Nrd1 from the nucleus delays expression of the *PHO5* gene. **(A)** Nothern blot analysis of total RNA from the parental Anchor Away (AA) and the corresponding Nrd1-AA strains upon addition of rapamycin to the growth medium. Nothern blots were probed specifically for sense and antisense *PHO5* transcripts, while *ACT1* RNA was used as a loading control. **(B)** Acid phosphatase induction kinetics in Nrd1-AA strain upon induction through phosphate starvation with (+Rap) or without addition of rapamycin (-Rap). Reported values represent the means and standard deviations of three independent experiments (n = 3). Indicated differences show the significant differences using an unpaired Student’s t test. Two (**) and four (****) asterisks denote a p-value lower than or equal to 0.01 and 0.0001, respectively. **(C)** Same as (B), but rapamycin was added one hour before induction.

### Transcription of *PHO5* AS RNA regulates *PHO5* gene expression *in cis*

To increase the transcription level of the *PHO5* AS transcript without using RNA degradation/termination mutant backgrounds, we inserted the strong constitutive *TEF1* promoter in the antisense configuration downstream of the *PHO5* gene ORF (Fig. 4A). We confirmed that this resulted in the *TEF1* promoter driving AS transcription at the *PHO5* gene locus by RT-qPCR, as the level of *PHO5* AS transcript in these mutant cells was ≈20 times higher than in the corresponding wild-type cells. Impressively, even under +P_i_ conditions in which the *PHO5* mRNA is only basaly expressed, *TEF1*-induced overexpression of the AS transcript caused a severalfold decrease in *PHO5* mRNA level, confirming the negative correlation between *PHO5* AS and mRNA transcript levels. What is more, Pho5 expression in these cells was delayed compared with wild-type cells and did not reach full expression level even after overnight induction (Fig. 4B). This result indicates that an artificially induced high constitutive level of AS transcription at the *PHO5* locus drives repression of the *PHO5* gene even after prolonged induction.

**Figure 4.**
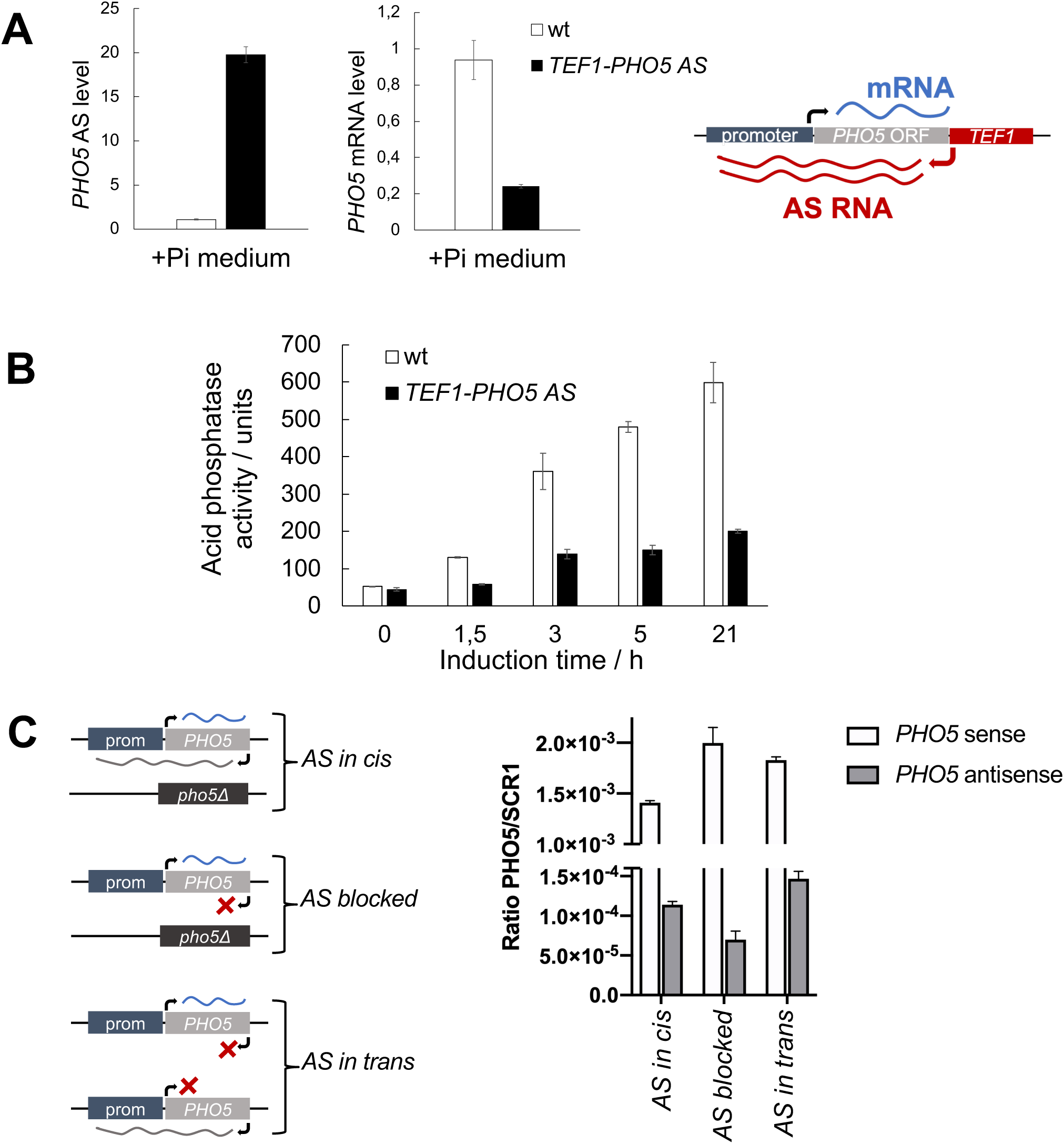
AS transcription represses the *PHO5* gene *in cis*. **(A)** Levels of *PHO5* AS and mRNA transcripts in the BMA41 wild-type and the corresponding *TEF1-PHO5 AS* strain at +P_i_ conditions, monitored by RT-qPCR. Values were normalized to *ACT1* RNA. Right: Scheme of the *PHO5* gene locus in the *TEF1-PHO5 AS* strain. **(B)** Acid phosphatase induction kinetics upon induction through phosphate starvation in wild-type BMA41 (wt) and corresponding *TEF1-PHO5 AS* mutant cells. Reported values represent the means and standard deviations of three independent experiments (n = 3). **(C)** Left: Scheme showing the *PHO5* gene locus in diploid strains in which *PHO5* AS is transcribed *in cis*, *in trans* or its transcription is blocked. Right: Levels of *PHO5* AS and S transcripts monitored by RT-qPCR in these strains. Values were normalized to *SCR1* RNA. Reported values represent the means and standard deviations of two independent experiments (n = 2).

Furthermore, we tested whether the *PHO5* AS transcript can regulate *PHO5* gene expression when expressed *in trans*, *i.e.* whether the AS transcript itself has a regulatory function. We constructed diploid strains (as in (9)) in which only one copy of the *PHO5* AS transcript was expressed either *in cis* (from the same chromosome as *PHO5* mRNA), *in trans* (from the opposite chromosome) and another one in which AS transcription in *cis* was blocked (Fig. 4C). Insertion of a terminator sequence to block AS transcription *in cis* resulted in only partial downregulation of *PHO5* AS level as shown by RT-qPCR. However, there was a marked increase in *PHO5* mRNA level in this diploid strain compared with the strain with native *PHO5* AS levels expressed *in cis* (Fig. 4C). Crucially, when *PHO5* AS was expressed *in trans* in addition to downregulation of its level *in cis, PHO5* S expression was higher than for the native locus indicating no repressive effect of the AS. These results argue that the act of AS transcription, rather than the AS RNA transcript itself, represses transcription of the *PHO5* gene.

### Block of AS transcription through dCas9 enhances the kinetics of *PHO5* gene expression

Given that accumulation of the *PHO5* AS transcript negatively affects *PHO5* gene transcription kinetics, blocking AS transcript production should enhance it. To specifically target *PHO5* AS transcription, we undertook a CRISPRi approach in which a catalytically dead Cas9 protein (dCas9) is directed by a guide RNA (gRNA) to interfere with AS transcription at the *PHO5* gene locus. The CRISPRi system blocks transcription due to physical collision between the elongating RNA Polymerase and the dCas9:gRNA complex (51). Furthermore, this system was shown to function in a strand-specific manner, by blocking transcription only when the nontemplate DNA strand of a transcription unit is targeted (51, 52). Therefore, we targeted dCas9 to the nontemplate strand of the AS transcription unit at the *PHO5* gene locus to block only AS transcription. First, we confirmed the presence of the dCas9 protein at the *PHO5* ORF by anti-Cas9 chromatin immunoprecipitation (ChIP). Notably, a strong peak of dCas9 binding at the *PHO5* gene ORF compared to a control strain not expressing the gRNA was observed (Fig. 5A), while no dCas9 binding could be detected at the *PHO5* promoter region covered by nucleosomes -4 and -1 (Fig. 5A). RNA levels in the Nrd1-AA strain with the active CRISPRi system were monitored by RT-qPCR and showed a highly reproducible decrease in *PHO5* AS levels compared to the control strain (Fig. 5B). This decrease was significant at the *PHO5* promoter and ORF regions without rapamycin addition or with rapamycin (*i.e.*, depletion of Nrd1 which induces AS transcription). These results are consistent with a dCas9-mediated transcriptional roadblock of AS transcription at the *PHO5* gene locus. After the addition of rapamycin, *PHO5* AS levels were increased in both the CRISPRi Nrd1-AA strain and the control Nrd1-AA strain. However, its levels in the CRISPRi strain remained significantly lower, maintaining the difference in levels already observed without the addition of rapamycin (Fig. 5B). These results demonstrated that the dCas9-mediated roadblock of AS transcription at the *PHO5* gene locus is robust and maintained after global induction of AS transcription, although AS transcription was not completely abolished.

**Figure 5.**
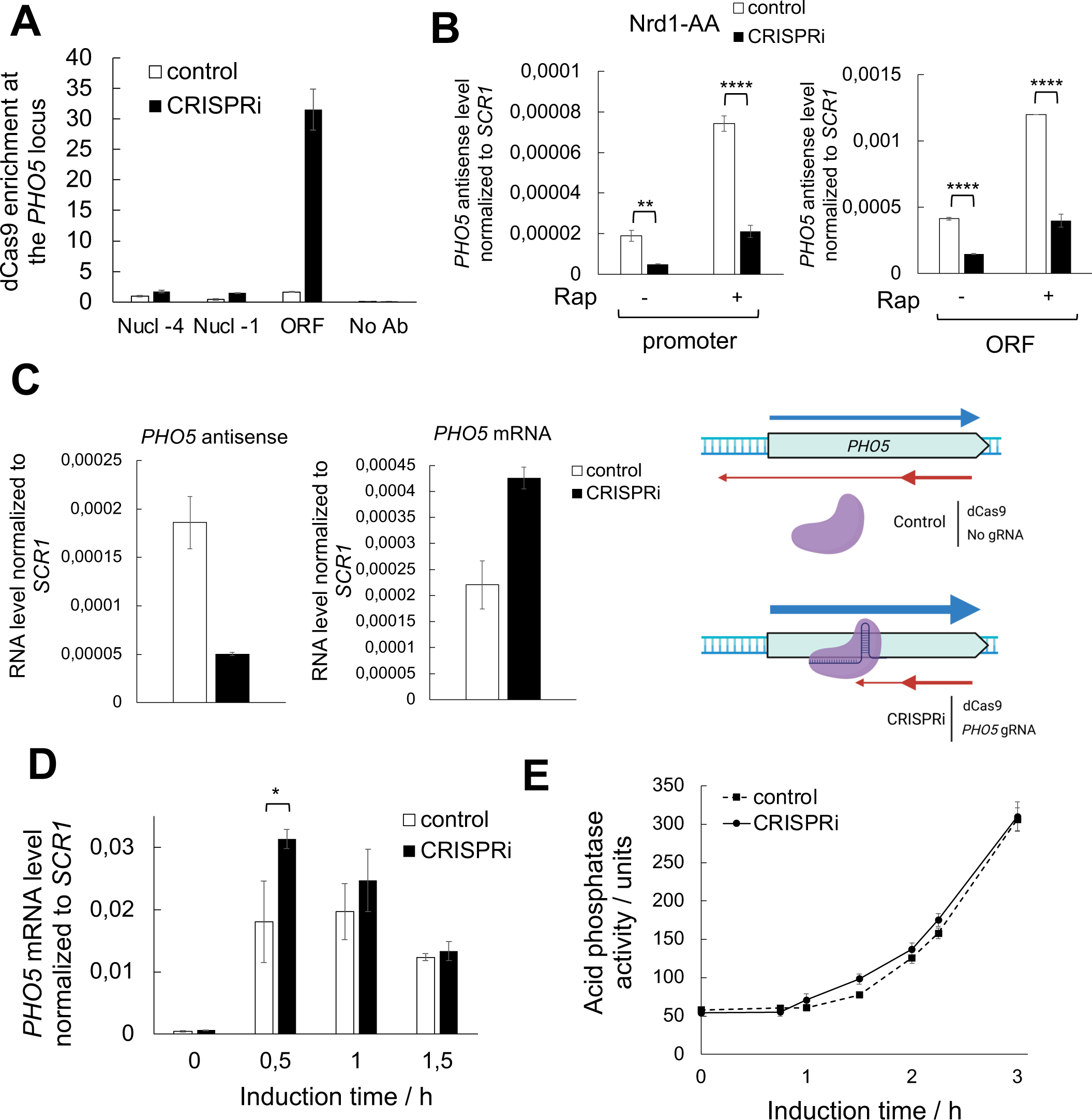
Targeting dCas9 to specifically block *PHO5* AS transcription enhances expression kinetics of the *PHO5* gene. **(A)** Chromatin immunoprecipitation (ChIP) analysis of dCas9 binding at the *PHO5* gene locus. Immunoprecipitated DNA was quantified by qPCR with primers specific for different regions of the *PHO5* promoter (Nucleosomes -4 and -1) and ORF regions. Both strains were transformed with a dCas9 expressing plasmid, while the CRISPRi strain was additionaly transformed with a plasmid expressing a gRNA targeted to strand-specifically block *PHO5* AS transcription and the control strain with the corresponding empty plasmid. Nucl - nucleosome, No Ab - no antibody ChIP control. **(B)** Levels of *PHO5* AS transcribed at the *PHO5* promoter and ORF regions at 0 h of induction in Nrd1-AA strain with or without addition of rapamycin (for 1 hour; to deplete Nrd1 and induce AS transcription) and an active CRISPRi system. RT-qPCR values were normalized to *SCR1* RNA. Reported values represent the means and standard deviations of three independent experiments (n = 3). Indicated differences show the significant differences using an unpaired Student’s t test. Two (**) and four (****) asterisks denote a p-value lower than or equal to 0.01 and 0.0001, respectively. **(C)** Levels of *PHO5* AS and mRNA transcripts in the CRISPRi and the corresponding control strain, monitored by RT-qPCR at 0 h of induction as in (B). Strains are Nrd1-AA with the absence of rapamycin (wild-type equivalent). Right: Scheme of the CRISPRi strategy used to block *PHO5* AS transcription. Strains were transformed with two plasmids, one expressing dCas9, and the other expressing or not a gRNA targeting the non-template strand of the *PHO5* AS transcription unit. **(D)** Levels of *PHO5* mRNA in the CRISPRi and the corresponding control strain upon induction through phosphate starvation monitored by RT-qPCR as in (B). Strains are same as in (C). Reported values represent the means and standard deviations of three independent experiments (n = 3). Indicated differences show the significant differences using an unpaired Student’s t test. One (*) and two (**) asterisks denote a p-value lower than or equal to 0.05 and 0.01, respectively. **(E)** Same as (D), but acid phosphatase induction kinetics were monitored by measuring acid phosphatase activity with whole cells.

Importantly, impairment of *PHO5* AS RNA elongation led to an increase in *PHO5* mRNA levels (Fig. 5C), clearly demonstrating the direct role of AS transcription in *PHO5* gene repression. Also, it argues in favour that the CRISPRi system strand-specifically blocked only AS transcription without significantly impacting mRNA transcription. We further tested if impairment of AS transcription with use of the CRISPRi system, would result in enhanced kinetics of *PHO5* gene expression. As expected, the kinetics of *PHO5* gene expression upon gene induction, were slightly faster when AS transcript production was impaired by dCas9 than in the control strain (Figs. 5D and 5E). This effect was noticed only at very early timepoints of gene induction (30 min for mRNA levels and 1,5 h for acid phospatase levels), possibly due to the dCas9 protein losing its roadblock function past a certain level of ongoing transcription.

## AS RNA elongation affects *PHO5* promoter chromatin structure

Since transcriptional activation of the *PHO5* promoter requires a large transition of its chromatin structure, we investigated whether the kinetics of *PHO5* promoter chromatin opening upon gene induction also inversely correlate with *PHO5* AS transcription. To this end, we examined the chromatin structure at the *PHO5* promoter with anti-histone H3 ChIP at nucleosome -2, which covers the high-affinity Pho4 binding site and is considered the critical nucleosome for *PHO5* chromatin remodeling (18). A higher histone occupancy was observed in *rrp6Δ* compared to wild-type cells already under repressive conditions (Fig. 6A). Accordingly, histone removal from the *PHO5* promoter was slower in *rrp6Δ* than in wild-type cells during the first hours of gene induction and reached a similar final level after 5 hours (Fig. 6A). To confirm the delayed kinetics of chromatin opening in *rrp6Δ* cells, we took advantage of the ClaI restriction enzyme accessibility assay, which quantifies the efficiency of cleavage by ClaI enzyme at nucleosome -2 of the *PHO5* promoter (Fig. 6B). Consistent with the anti-histone H3 ChIP, the accessibility of the ClaI site at the *PHO5* promoter was lower in *rrp6Δ* and *air1Δair2Δ* than in wild-type cells during the first hours of gene induction (Fig. 6C). These results show that AS transcription mediates a negative effect on *PHO5* transcriptional activation by influencing the chromatin structure at its promoter region.

**Figure 6.**
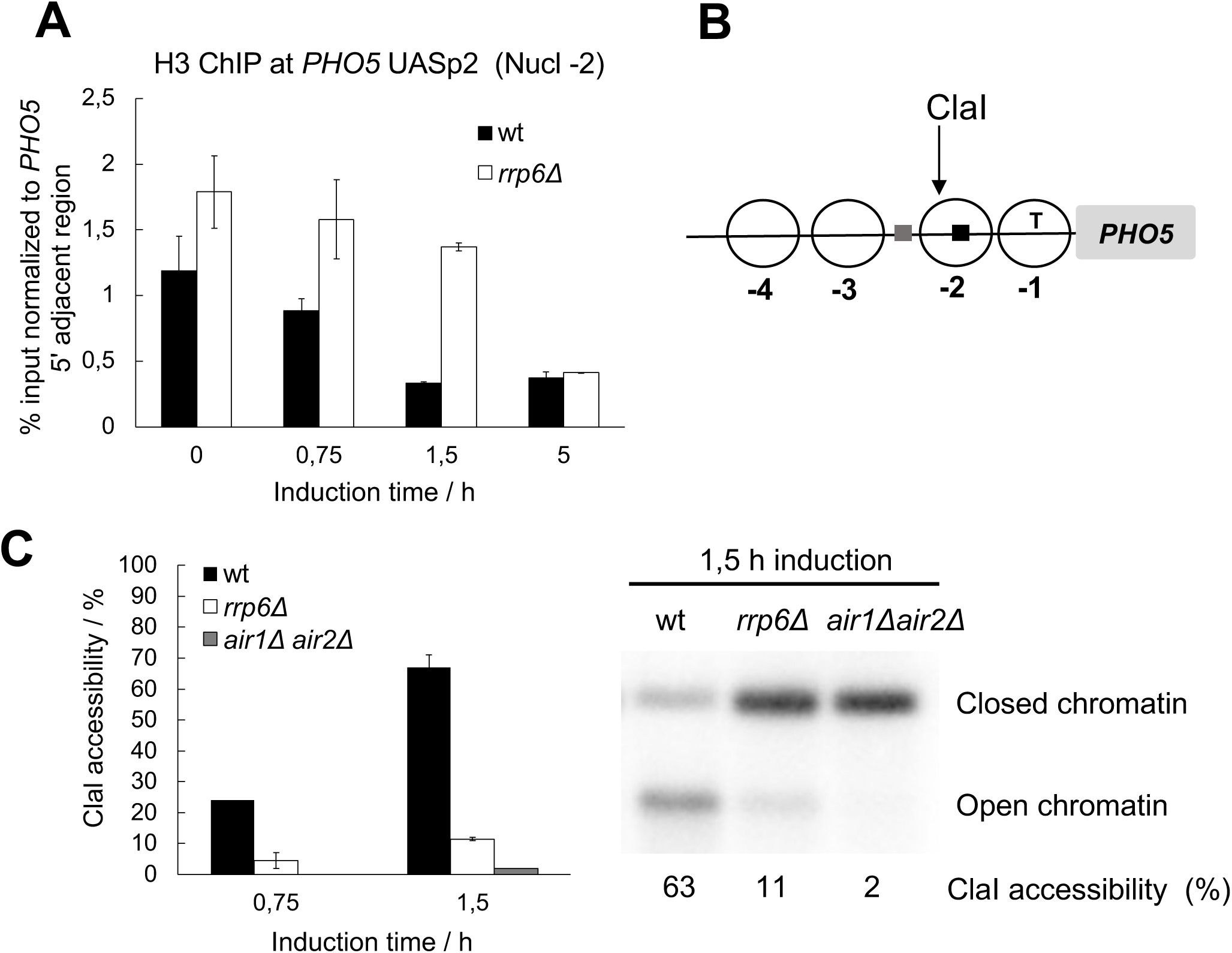
*PHO5* AS elongation negatively affects kinetics of histone removal at the *PHO5* gene promoter upon induction. **(A)** ChIP analysis of histone H3 binding at nucleosome -2 of the *PHO5* gene promoter in wild-type BMA41 (wt) and corresponding *rrp6Δ* cells upon induction through phosphate starvation. Immunoprecipitated DNA was quantified by qPCR and normalized to a control genomic region adjacent to the *PHO5* gene locus. **(B)** Scheme of the *PHO5* gene promoter region. Nucleosomes are denoted by circles, Pho4 binding sites by squares (gray - low affinity, black - high affinity) and the TATA box by the letter T. Site of cleavage with the ClaI restriction enzyme is denoted by a black arrow. **(C)** Kinetics of *PHO5* promoter opening monitored by ClaI accessibility at nucleosome -2 after induction as in (A).

Our results suggest that AS transcription at the *PHO5* gene locus locks the chromatin structure of the *PHO5* promoter in a more repressive configuration that is harder to remodel (Fig. 6). This could be due to the activity of HDACs, which have been shown to negatively affect chromatin structure at the *PHO5* promoter (20, 21). Remarkably, inactivation of the HDAC Rpd3 in the *rrp6Δ* mutant background does not affect the level of the *PHO5* AS RNA, but it restores transcription activation of the *PHO5* gene to the level or even higher than in wild-type cells, as shown by tiling arrays and RT-qPCR with single and double deletion mutant cells ((14); Fig. 7A). Accordingly, the expression kinetics of acid phosphatase measured with the *rpd3Δ rrp6Δ* double mutant cells are not delayed compared to wild-type cells, in contrast to the corresponding *rrp6Δ* single mutant cells (Fig. 7B). Consistent with this, expressing the *PHO5* AS-blocking CRISPRi system leads to faster gene expression kinetics in wild-type and *rrp6Δ*, but not in *rpd3Δ* and *rpd3Δ rrp6Δ* double mutant cells (Fig. S3). These results demonstrate that the *PHO5* AS transcript acts *via* a pathway that involves histone deacetylation. Gcn5, the catalytic subunit of the SAGA and ADA complexes, is known to be the major histone acetyltransferase that enables physiological gene induction kinetics at the *PHO5* promoter (23, 24). We reasoned that in the absence of Gcn5, *i.e.* when the majority of histone acetylation normally present at the *PHO5* gene promoter is reduced, an *rrp6Δ* strain should have no additional effect on the kinetics of *PHO5* gene expression. Indeed, the kinetics of acid phosphatase expression in *gcn5Δrrp6Δ* double mutant strain are the same as in the *gcn5Δ* single mutant throughout the induction period (Fig. 7C). Taken together, these results support that AS transcription-mediated repression of *PHO5* gene expression occurs *via* histone deacetylation.

**Figure 7.**
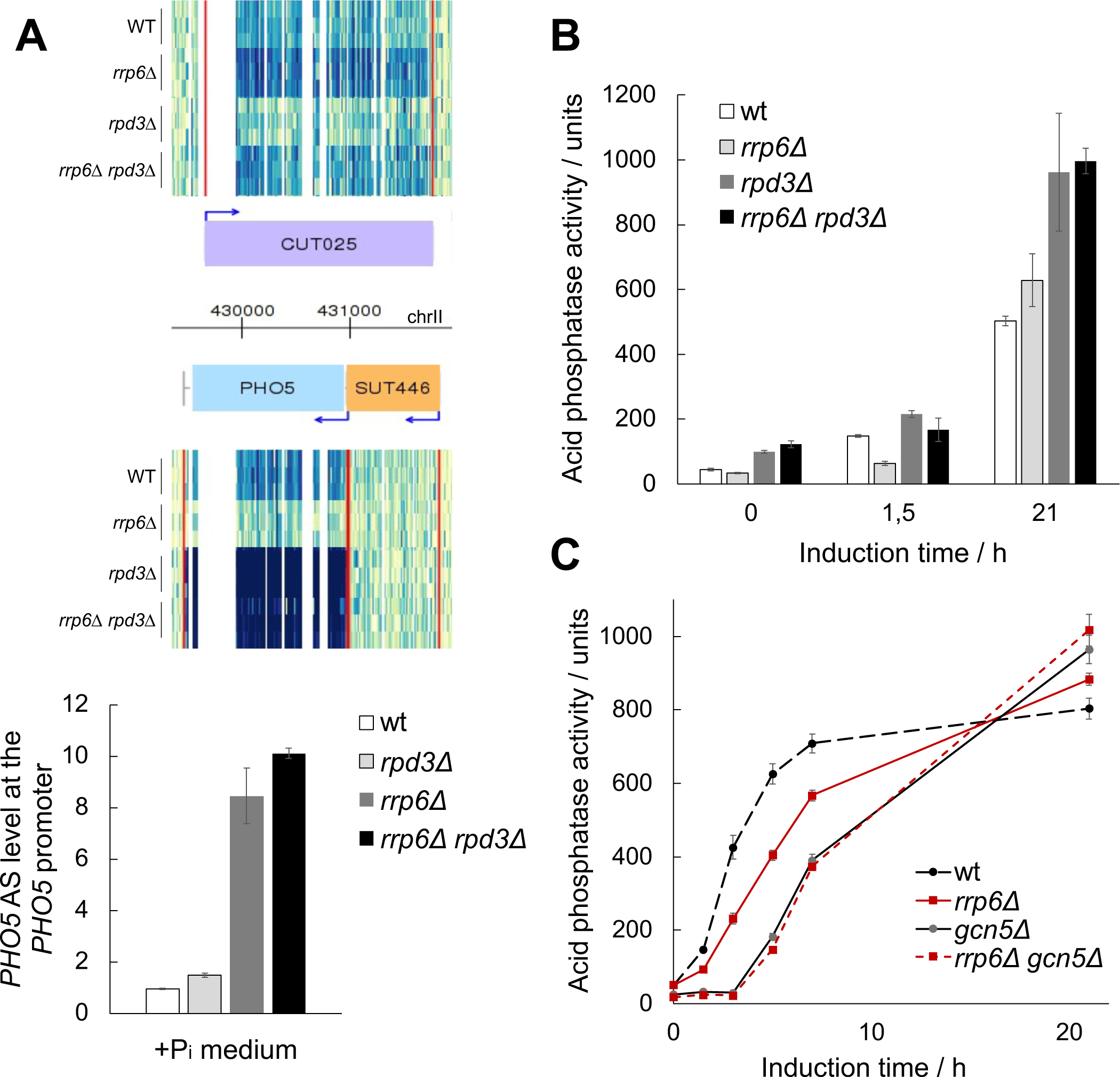
*PHO5* AS elongation affects *PHO5* gene expression *via* histone acetylation. **(A)** Heatmap of the *PHO5* gene locus in wild-type W303 (wt), *rpd3Δ, rrp6Δ*, and *rrp6Δrpd3Δ* mutant cells. Snapshot of tilling arrays intensities from (14) at the *PHO5* locus for the Watson (W, upper half) and the Crick (C, lower half) strands. Three replicates of each strain are represented. A darker signal depicts a higher score of RNA expression. The red vertical lines represent the inferred coding and non-coding genes boundaries. Below: Levels of *PHO5* AS transcript measured by RT-qPCR with the same strains at +Pi conditions. Values were normalized to *ACT1* RNA. **(B)** Acid phosphatase induction kinetics in wild-type W303 (wt) and the corresponding *rrp6Δ*, *rpd3Δ* and *rrp6Δrpd3Δ* cells upon induction through phosphate starvation. Reported values represent the means and standard deviations of three independent experiments (n = 3). **(C)** As in (B), but for wild-type BY4741 (wt) and the corresponding *rrp6Δ*, *gcn5Δ* and *rrp6Δgcn5Δ* mutant cells.

### AS transcription negatively affects recruitment of RSC to the *PHO5* promoter

Histone acetylation plays two important roles in transcriptional activation. It neutralizes the positive charge of lysine groups, thereby weakening histone-DNA interactions, and it also provides docking sites for the bromodomains of proteins involved in transcriptional regulation. RSC (Remodels Structure of Chromatin) complex is the most abundant and the only essential remodeler in yeast and contains seven of the fourteen bromodomains identified in *S. cerevisiae* (26, 53). RSC was found to be the major remodeler among the five chromatin remodelers involved in the chromatin remodeling process at the *PHO5* promoter (29). Its partial depletion, achieved by a temperature-sensitive mutant of its catalytic subunit *sth1^td^*, resulted in a strong delay in promoter chromatin structure opening and, consequently, delayed kinetics of acid phosphatase expression upon *PHO5* gene induction.

To first test the hypothesis that histone acetylation recruits RSC to the *PHO5* gene promoter upon induction, we used the anchor away system to deplete its catalytic subunit Sth1 from the nucleus in Sth1-AA and corresponding *gcn5Δ* mutant cells. Because Sth1 is essential for cell viability, we first attempted to induce its depletion in parallel with the induction of the *PHO5* gene. Addition of rapamycin upon shifting the cells to phosphate-free medium caused a delay in acid phosphatase expression kinetics similar to the partial depletion through *sth1^td^* (Fig. 8A). In *gcn5Δ* mutant cells, this partial depletion leads to an additive effect on acid phosphatase expression kinetics. However, since RSC is very abundant, it is possible that the partial depletion of Sth1 still leaves a lot of active RSC complex in the nucleus in the first hours of gene induction. We therefore added rapamycin two hours before *PHO5* gene induction to achieve more extensive RSC depletion before shifting the cells to phosphate-free medium. Addition of rapamycin two hours before gene induction resulted in an epistatic effect of the Sth1 depletion. Upon simultaneous inactivaton of Gcn5 and RSC, acid phosphatase expression kinetics were severely delayed, but reached overnight levels comparable to wild-type (Sth1-AA -Rap) cells (Fig. 8A). This result positions Gcn5 and RSC in the same pathway of *PHO5* gene transcriptional activation and speaks in favour of a link between RSC recruitment and Gcn5-mediated acetylation upon induction.

**Figure 8.**
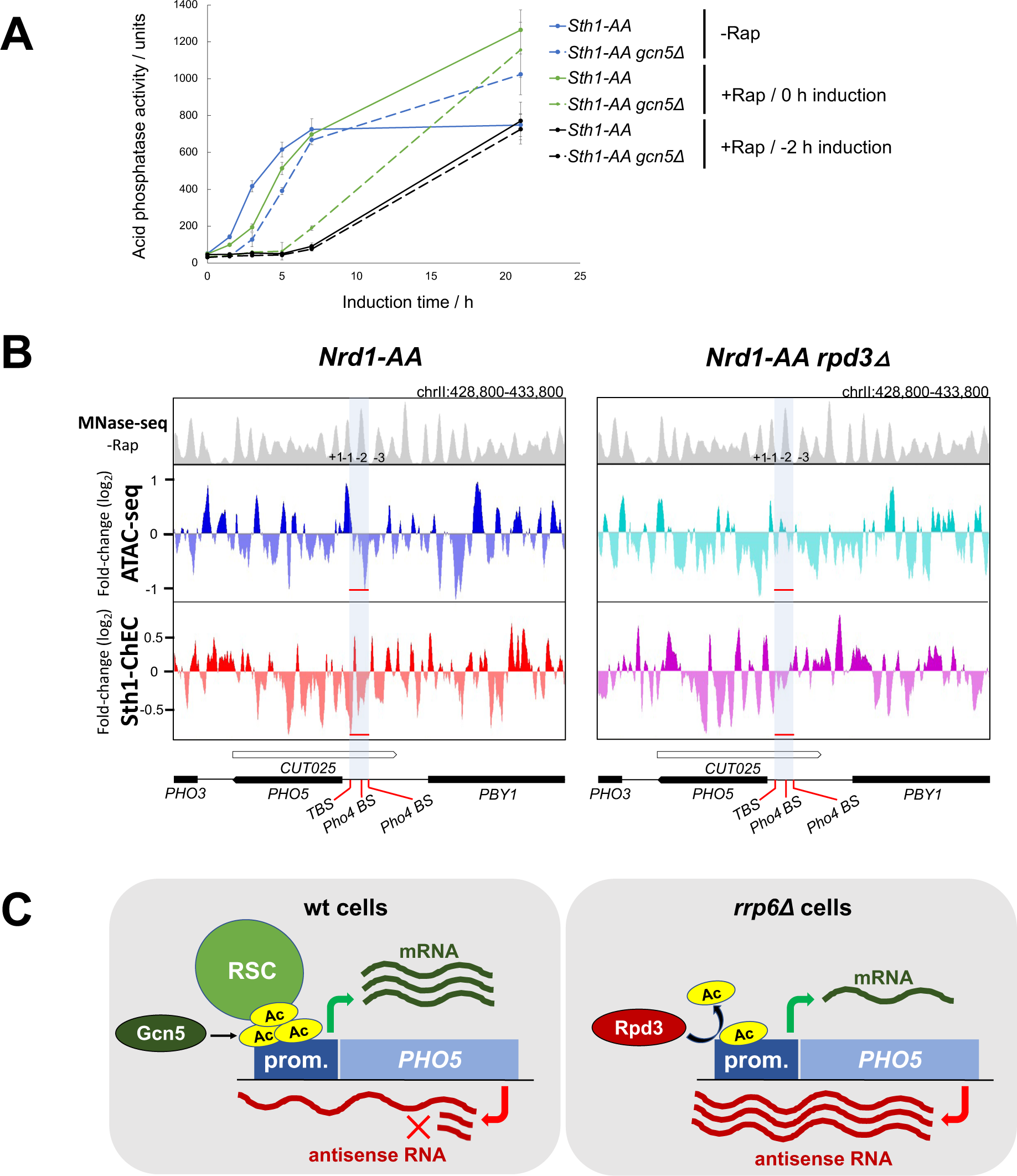
Chromatin remodeling at the *PHO5* gene promoter is negatively affected by *PHO5* AS elongation. **(A)** Acid phosphatase induction kinetics in Sth1-AA and the corresponding *gcn5Δ* cells upon induction through phosphate starvation without (-Rap) or with addition of rapamycin (+Rap) at indicated times. Reported values represent the means and standard deviations of three independent experiments (n = 3). **(B)** Snapshot of the *PHO5* gene locus in Nrd1-AA and the corresponding *rpd3Δ* strain from MNase-seq (-Rap), ATAC-seq (+Rap/-Rap) and Sth1-ChEC (+Rap/-Rap) experiments. Data is from (12). **(C)** Proposed model for how AS RNA regulates transcription of the *PHO5* gene *via* remodeling of promoter chromatin structure. In wild-type cells, antisense RNA transcription is terminated by the NNS complex and degraded by the RNA exosome. Histones at the *PHO5* gene promoter are acetylated by Gcn5 and serve as docking sites for recruitment of the chromatin remodeling complex RSC, thus enabling physiological kinetics of promoter opening and gene induction. In *rrp6Δ* cells, read-through of the AS transcript into the *PHO5* promoter region results in increased recruitment of the histone deacetylase Rpd3 and subsequently in hypoacetylation and decreased recruitment of RSC. This results in delayed kinetics of promoter opening and induction of the *PHO5* gene.

To test the effect of Sth1-AA depletion in *rrp6Δ* mutant cells, we monitored acid phosphatase expression kinetics upon addition of rapamycin (Fig. S4). Even when rapamycin was added two hours before induction to achieve more complete inactivation of RSC, it resulted in an additive effect on acid phosphatase expression kinetics with the *rrp6Δ* mutation. It is possible that Rrp6 and RSC regulate *PHO5* gene expression through at least partially independent pathways. However, because these cells barely induced the *PHO*5 gene, as indicated by the levels of acid phosphatase activity measured after overnight induction, we cannot rule out the possibility that this additive effect is due to the severely impaired cell viability because of the Sth1-AA depletion in the slow-growing *rrp6Δ* background.

To directly answer the question of whether AS-induced deacetylation of the *PHO5* promoter may inhibit the recruitment of RSC, resulting in a more closed chromatin conformation, we took advantage of our genomic analyses recently performed with Nrd1-AA and Nrd1-AA *rpd3Δ* cells with and without the addition of rapamycin (12). We examined the *PHO5* gene locus in the Micrococcal Nuclease sequencing (MNase-seq), Assay for Transposase-Accessible Chromatin using sequencing (ATAC-seq) and Sth1-Chromatin Endogenous Cleavage-sequencing (Sth1-ChEC-seq) datasets, which give us information about the chromatin conformation and Sth1 binding at the *PHO5* promoter upon induction of AS transcription (+Rap/-Rap) and depending on the presence of Rpd3 (Fig. 8B). The ChEC-seq data show the fold change in association of Sth1, the ATP-ase subunit of RSC, with chromatin upon induction of AS transcription (+Rap/-Rap). In addition, the ATAC-seq data provide us with information about chromatin accessibility under the same conditions. In Nrd1-AA cells there is a negative fold change, *i.e.* a decrease in Sth1 binding, associated with a decrease in chromatin accessibility upon addition of rapamycin, for the region encompassing nucleosome -2 of the *PHO5* promoter (the position of which was determined using MNase-seq data in -Rap) (Fig. 8B). Conversely, in isogenic *rpd3Δ* cells, addition of rapamycin has a much smaller effect on Sth1 binding or chromatin accessibility in this region (Fig. 8B). When comparing the two biological experimental replicates, the log2 values for the change Sth1 binding (+Rap/-Rap) were consistently lower in Nrd1-AA compared to isogenic *rpd3Δ* cells (-0,3947 and -0,566 *vs.* -0,2057 and -0,2841, respectively, calculated over the middle 40 bp region of nucleosome -2). These data argue in favour of a model in which read-through of AS transcription acts *via* recruitment of histone deacetylases to the *PHO5* gene promoter, the activity of which results in decreased recruitment of the RSC complex (Fig. 8C).

## Discussion

The role of non-coding RNAs in regulation of gene expression could not be appreciated until recent advances in high-throughput methods facilitated their detection and characterization. From a gene-centered view, non-coding RNAs can be transcribed in tandem with genes, *i.e.* from the same strand as the gene, or from the opposite strand, resulting in production of antisense (AS) non-coding RNAs. Apart from a few isolated examples, production of AS non-coding RNAs is generally thought to have a repressive *cis*-regulatory effect on the expression of associated mRNAs (6, 8, 54). This seems to be particularly the case when transcription of AS non-coding RNAs invades promoters of coding genes (9, 14, 55). In light of this current view, we felt compelled to reexamine the role of AS transcription at the model yeast *PHO5* gene locus, which was originally suggested to support gene activation (33). In this work, we show a clear negative role for AS transcription in *PHO5* gene expression. By leveraging mutant backgrounds in which AS transcription is constitutively enhanced or inducible and artificially driving its expression from a strong promoter *in cis*, we show that increased *PHO5* AS elongation frequency correlates with decreased expression of the corresponding mRNA. Furthermore, we demonstrate that the use of a CRISPRi system that specifically blocks AS transcription at the *PHO5* gene locus increases the level of *PHO5* mRNA and enhances its induction kinetics upon phosphate depletion. Importantly, these observations show that AS RNA transcription has an impact on *PHO5* gene expression in wild-type cells, and not only upon enhanced AS RNA stabilisation in strains mutant for RNA degradation factors. We also show that AS RNA transcription regulates expression of the *PHO5* gene only when transcribed *in cis*, and not *in trans*. The role of *PHO5* AS transcription is therefore reminiscent of the role of AS transcription in maintaining the tight repression of quiescence-related transcripts during the exponential growth phase, recently demonstrated by Nevers et al. (9). A previous study suggesting a positive regulatory role for *PHO5* AS transcription achieved AS inactivation by incorporating a full-length marker gene sequence with its promoter region in the middle of the *PHO5* gene ORF (33). This major perturbation of the *PHO5* gene locus may have resulted in experimental artefacts, highlighting the need for precise interventions, such as those achieved by the CRISPRi system, to perform functional analyses of AS transcripts (56).

There are now several well-described examples of yeast gene loci at which either antisense or upstream non-coding transcription that extends through a coding sense promoter has an inhibitory effect on its transcription initiation (13, 57–60). In most cases, it is likely that elongation of non-coding transcription leads directly to displacement of transcription factors (TFs) and/or the preinitiation complex (PIC) or that the recruitment of TFs or the PIC to these gene promoters is decreased as a consequence of a more repressive chromatin configuration established at the promoter region due to elongation of non-coding transcription (see (61) for a review). This model is supported by whole-genome analyses showing that invasion of gene promoters by AS transcription leads to increased histone occupancy and altered recruitment of chromatin-modifying and -remodeling complexes (10, 12, 62). At the tandemly transcribed *SRG1* lncRNA/*SER3* protein-coding gene locus, non-coding transcription has been shown to cause nucleosome deposition at the gene promoter, thereby repressing *SER3* transcription (63). As another example, we have shown that AS transcription at the *PHO84* gene locus silences the corresponding gene by recruiting HDACs to its promoter region (13). The AS RNA does not recruit the HDACs directly, but the act of its transcription promotes a histone methylation-based mechanism to restore the repressive chromatin structure in the wake of the elongating RNA Pol II. The histone methyltransferase Set2 associates with the elongating RNA Pol II and catalyses H3K36 methylation, a mark read by the Eaf3 chromodomain of the HDAC Rpd3 (64). Consistent with this, our recent genome-wide study in yeast has shown that AS transcription leads to deacetylation of a subpopulation of -1/+1 nucleosomes associated with increased H3K36 methylation, which in turn leads to decreased binding of the RSC chromatin-remodeling complex and sliding of nucleosomes into previously nucleosome-depleted regions (12). We have now shown that elongation of *PHO5* AS under repressive conditions leads to increased histone occupancy at the *PHO5* gene promoter and slower histone removal upon gene induction. Moreover, the negative effect of AS RNA elongation on *PHO5* gene activation is mitigated by inactivation of Rpd3, suggesting a histone acetylation-based regulatory mechanism that may affect the recruitment of RSC, a chromatin remodeler that plays an important role in *PHO5* gene promoter opening (29). This is supported by ChEC-seq of Sth1, the catalytic subunit of RSC, showing a decrease in its recruitment to the *PHO5* gene promoter upon induction of antisense transcription, that is suppressed by inactivation of Rpd3.

*PHO5* belongs to a group of ∼100 genes that are more transcribed in AS direction as a non-coding transcript than in the sense orientation as an mRNA in a standard medium (Fig. 1B). In such culture conditions, the Pho4 transcriptional activator is rarely located in the nucleus (18). Thus, as we proposed in (44) for the SAGA-dependent gene class to which *PHO5* belongs, the steady-state chromatin structure of the promoter NDR might be maintained tightly closed by ongoing AS transcription. What may also be relevant to this mechanism is the recently discovered autoregulatory mechanism of the SAGA complex, which is induced in response to environmental changes such as phosphate starvation conditions (65). The SAGA catalytic subunit Gcn5 has been shown to acetylate the Ada3 subunit, which promotes dimerization of the SAGA complex and in turn leads to higher efficiency of SAGA-catalysed histone acetylation. *PHO5* expression was shown to correlate negatively with decreasing levels of Ada3 acetylation and consequently lower efficiency of histone acetylation by Gcn5. The same was also found for *SUC2* transcription, which is induced during growth in sucrose-containing media. Of importance to our work is the finding that of the 8 known histone deacetylases, the Ada3 subunit is deacetylated only by Rpd3, but the mechanism of its recruitment to SAGA remains to be elucidated. Therefore, the enhanced recruitment of Rpd3 mediated by AS transcription may play a dual role in regulating *PHO5* gene expression, considering that Rpd3 deacetylates promoter histones and Gcn5, both of which contribute to transcriptional repression. It remains to be investigated whether such a regulatory mechanism of AS transcription-mediated repression could be a common mechanism for AS transcription-induced repression of stress-inducible and SAGA-dependent genes regulated by promoter chromatin structure remodeling.

The regulatory roles of non-coding RNAs are intertwined with that of chromatin structure. Not only does non-coding transcription affect chromatin structure, but chromatin structure also determines where and how often non-coding RNAs are transcribed. This fact is increasingly appreciated with respect to the directionality of transcription at promoters of coding genes. Specifically, chromatin modifiers such as the HDAC Hda1, and chromatin remodelers such as RSC, have been shown to dictate promoter directionality by attenuating divergent non-coding transcription (66, 67). Furthermore, chromatinization of DNA limits aberrant transcription that would otherwise occur on naked DNA, as was recently demonstrated through *in vitro* experiments by the Kornberg group (68). In this study, a chromatinized *PHO5* gene locus fragment was transcribed seven times more from the physiological transcription start site than the same naked DNA locus, and also resulted in transcription patterns more similar to those seen *in vivo*. Although only chromatin was considered in this study, it would be interesting to also investigate non-coding transcription using a similar *in vitro* transcription system.

Chromatin remodeling complexes and non-coding RNAs are important regulators of gene expression, and therefore dysregulation of either of these factors may affect the development and progression of various cancers. The SWI/SNF family of chromatin remodeling complexes includes the SWI/SNF complex with its catalytic subunits BRG1 or BRM in humans, and the SWI/SNF and RSC complex with their catalytic subunits Snf2 and Sth1, respectively, in yeast. Numerous associations between chromatin remodelers of this family and long non-coding RNAs have been detected in human cancers (reviewed in (69)). These complexes and the corresponding regulatory non-coding RNAs therefore represent promising diagnostic and therapeutic targets. Transcription of long non-coding RNAs is particularly important for the yeast genome, which has a very high gene density, such that many of them overlap coding gene ORFs or promoter regions. Another reason why budding yeast is a good model for studying the transcription of such long (≥200 nt) non-coding RNAs is that it exclusively synthesizes this non-coding transcript class since its divergence from other yeasts and the loss of the RNAi system that produces small non-coding RNAs (70). In addition, extensively studied gene loci, such as the yeast *PHO5* gene, are invaluable for mechanistic studies of gene regulation. Studies of the *PHO5* gene and its promoter region made an immense contribution to deciphering the mechanisms of gene regulation through chromatin remodeling (18) and our study now opens the possibility to focus on non-coding transcription in this system.

## Acknowledgements

I.S. and A.N. dedicate this paper to A. Rachid Rahmouni, who always provided support and encouragement. We are grateful to P. Korber for extensively commenting on the manuscript and providing help with the restriction nuclease accessibility assay.

---

**Figure S1.**
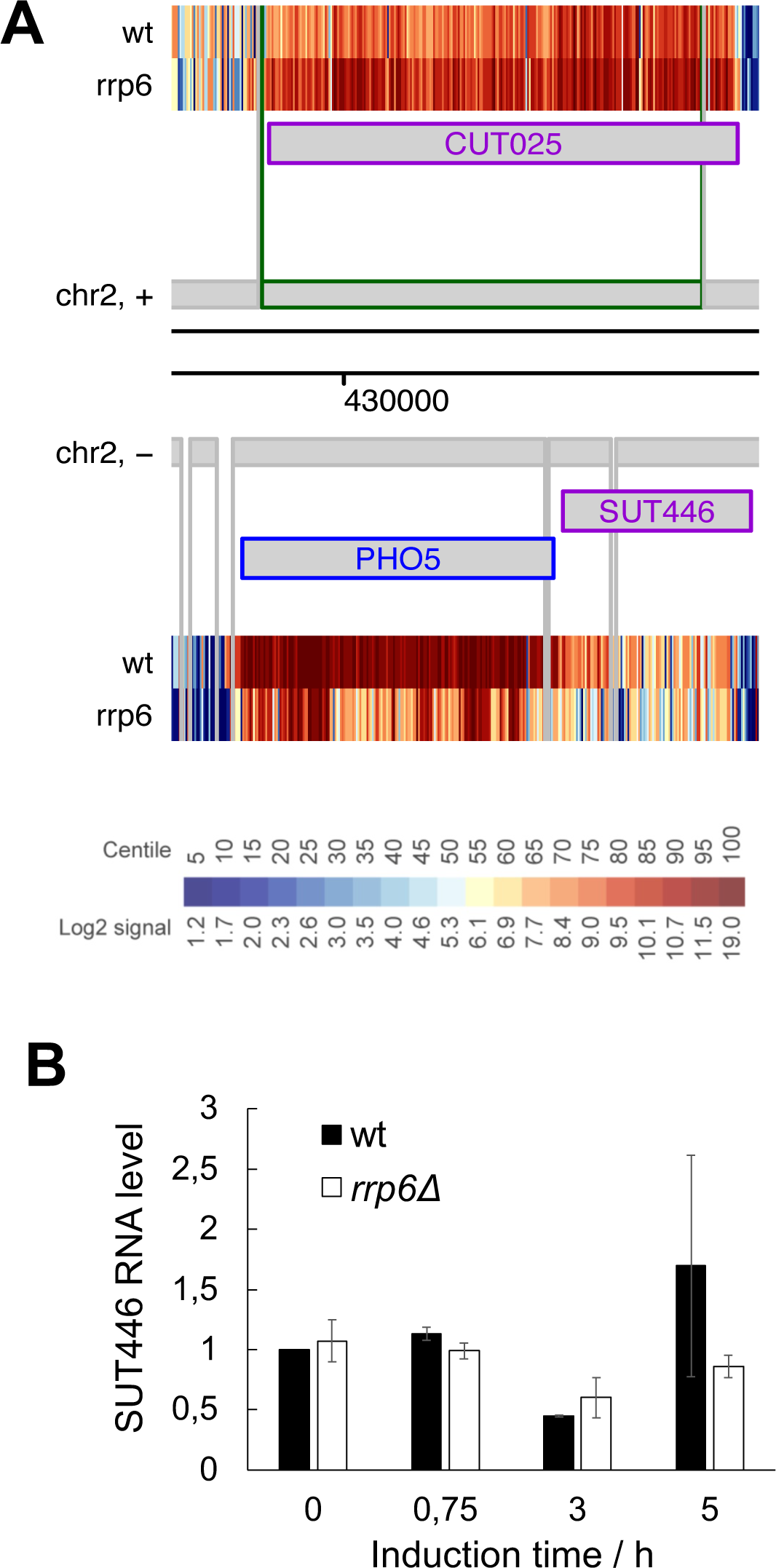
Non-coding transcripts CUT025 and SUT446 are transcribed at the *PHO5* gene locus. **(A)** A heatmap summarising tiling array expression data at the *PHO5* gene locus in wild-type W101 (wt) and corresponding *rrp6Δ* cells. Data is from (34) and is visualized with the SGV Genomics Viewer (71). **(B)** Levels of SUT446 in wild-type BMA41 (wt) and corresponding *rrp6Δ* mutant cells upon induction through phosphate starvation. RT-qPCR values were normalized to *PMA1* RNA and expressed relative to transcript abundance in wild-type cells at repressive conditions (0 h of induction), which was set to 1.

**Figure S2.**
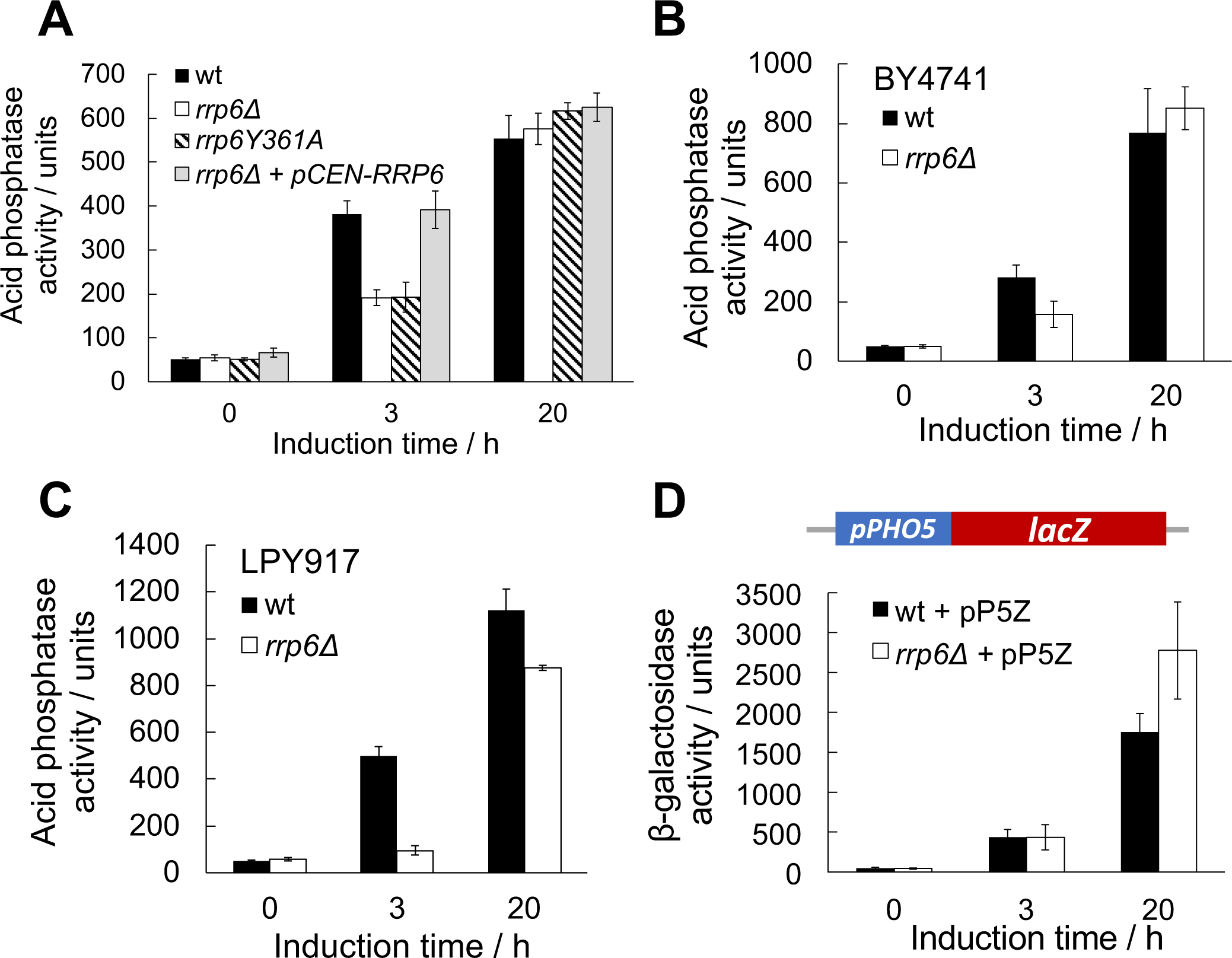
Delayed expression kinetics of the *PHO5*, but not the *lacZ* gene under regulation of the *PHO5* promoter in *rrp6Δ* mutant cells. **(A)** Acid phosphatase induction kinetics in wild-type BMA41 (wt) and corresponding mutant cells upon induction through phosphate starvation. The strain *rrp6Y361A* carries a point mutation at the *RRP6* genomic locus which abolishes exonuclease activity of Rrp6. Plasmid pCEN-RRP6 is a centromeric plasmid which carries the *RRP6* gene under regulation of its native promoter. Reported values represent the means and standard deviations of three independent experiments (n = 3). **(B)** Same as (A), but for wild type and corresponding *rrp6Δ* mutant cells from the BY4741 genetic background. **(C)** Same as (A), but for wild type and corresponding *rrp6Δ* mutant cells from the LPY917 genetic background. **(D)** Beta-galactosidase induction kinetics in wild-type BMA41 (wt) and corresponding *rrp6Δ* cells transformed with a reporter plasmid pP5Z carrying the *lacZ* gene under the control of the *PHO5* promoter upon induction through phosphate starvation. Reported values represent the means and standard deviations of three independent experiments (n = 3).

**Figure S3.**
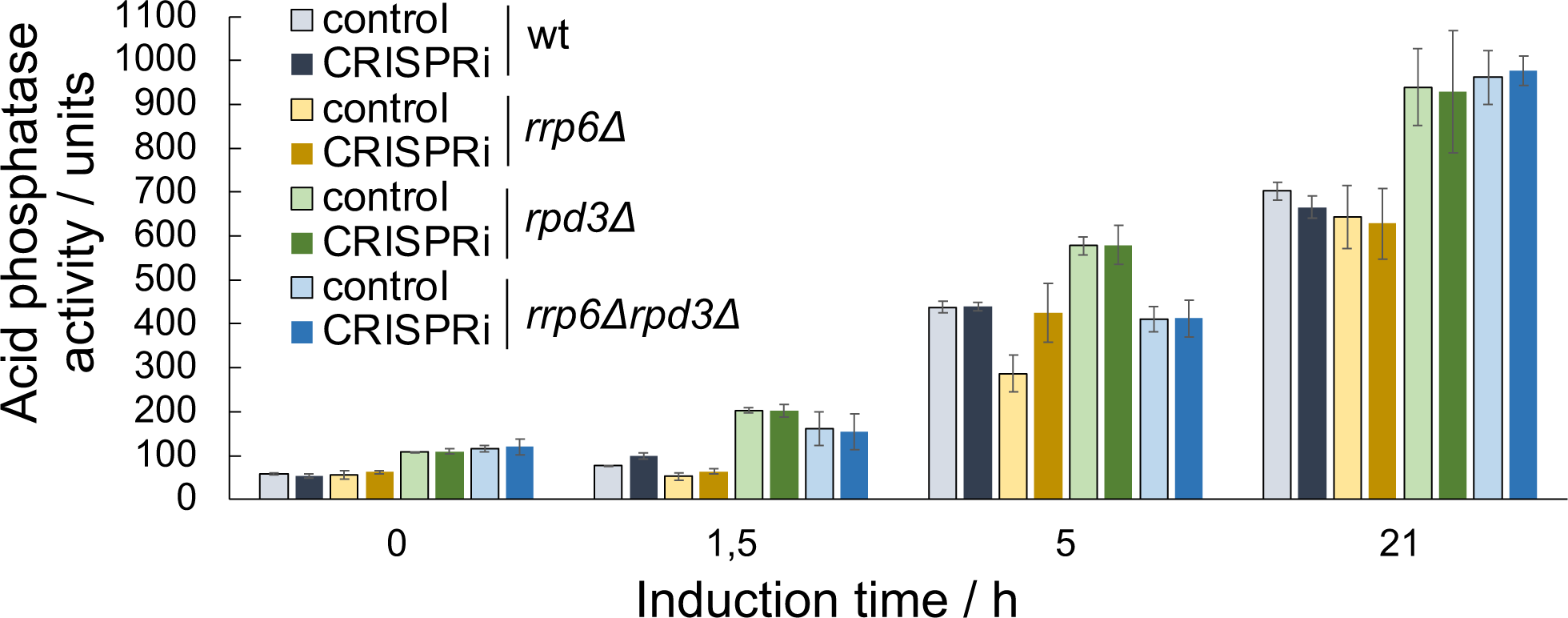
Expression of the *PHO5* AS-blocking CRISPRi system leads to faster gene expression kinetics in wt and *rrp6Δ*, but not in *rpd3Δ* and *rpd3Δ rrp6Δ* double mutant cells. Acid phosphatase induction kinetics in wild-type BMA41 (wt) and corresponding deletion mutant cells for Rrp6 and Rpd3, with and without expression of the CRISPRi system which blocks *PHO5* AS transcription, upon induction through phosphate starvation. Reported values represent the means and standard deviations of three independent experiments (n = 3).

**Figure S4.**
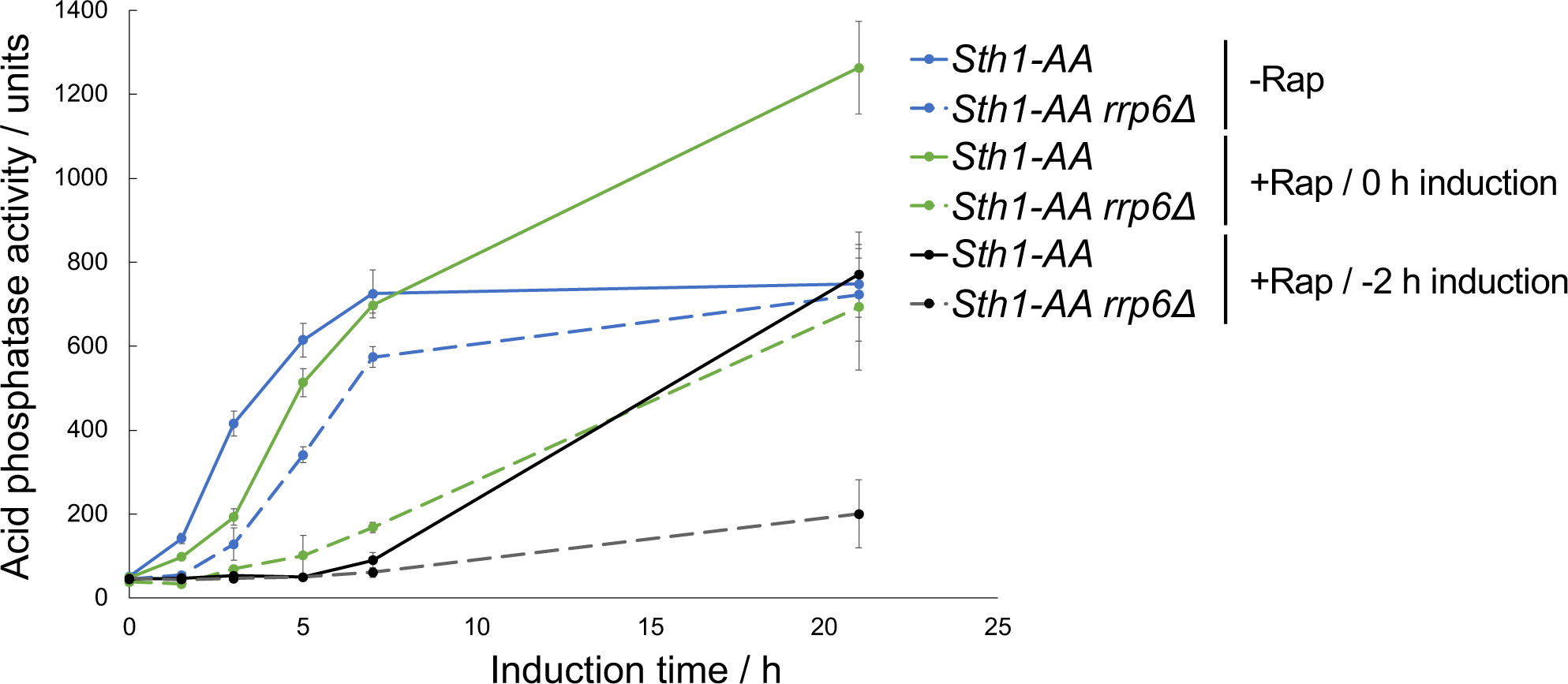
Effect of simultaneous inactivation of Sth1 and Rrp6 on *PHO5* gene expression. Acid phosphatase induction kinetics in Sth1-AA and the corresponding *rrp6Δ* cells upon induction through phosphate starvation without (-Rap) or with addition of rapamycin (+Rap) at indicated times. Reported values represent the means and standard deviations of three independent experiments (n = 3).

