## Supplemental tables for "Antisense non-coding transcription represses the *PHO5* model gene at the level of promoter chromatin structure"

**Table S1.** ***S. cerevisiae*** **strains**

| Strain ID | Genotype | Source |
|---|---|---|
| BMA41 wild-type | *MATa ade2-1 ura3-1 leu2-3,112 his3-11,15 trp1Δ can1-100* | (1) |
| BMA41 *rrp6Δ* | BMA41 with *rrp6Δ::KanMX4* | (2) |
| BMA41 Rrp6-Y361A | BMA41 with *rrp6Y361A* | (3) |
| BMA41 *rrp47Δ* | BMA41 with *rrp47Δ::KanMX4* | (3) |
| BMA41 *trf4Δ* | BMA41 with *trf4Δ::KanMX4* | (3) |
| BMA41 *trf5Δ* | BMA41 with *trf5Δ::KanMX4* | (3) |
| BMA41 *mpp6Δ* | BMA41 with *mpp6Δ::KanMX4* | (3) |
| BMA41 *air1Δ* | BMA41 with *air1Δ::KanMX4* | (3) |
| BMA41 *air2Δ* | BMA41 with *air2Δ::KanMX4* | (3) |
| BMA41 *air1Δ air2Δ* | *MATa ade2-1 ura3-1 leu2-3,112 his3-11,15 trp1Δ can1-100 air1Δ::HIS3 air2Δ::KanMX4* | (2) |
| BMA41 *TEF1-PHO5AS* | BMA41 with *TEF1-PHO5AS::KanMX4* | This work |
| *dis3Δ* + pDis3 | *MATa ade2-1 ura3-1 leu2-3,112 his3-11,15 trp1-1 can1-100 dis3Δ::KanMX4 [pBS3269-DIS3, LEU2]* | (3) |
| *dis3Δ* + pDis3-endo$^-$ | *MATa ade2-1 ura3-1 leu2-3,112 his3-11,15 trp1-1 can1-100 dis3Δ::KanMX4 [pBS3278-dis3D171N, LEU2]* | (3) |
| *dis3Δ* + pDis3-exo$^-$ | *MATa ade2-1 ura3-1 leu2-3,112 his3-11,15 trp1-1 can1-100 dis3Δ::KanMX4 [pBS3270-dis3D551N, LEU2]* | (3) |
| BY4741 wild-type | *MATa his3Δ1 leu2Δ0 met15Δ0 ura3Δ0* | (4) |
| BY4741 *rrp6Δ* | BY4741 with *rrp6Δ::KanMX4* | (5) |
| BY4741 *gcn5Δ* | BY4741 with *gcn5Δ::KanMX4* | EUROSCARF |
| BY4741 *gcn5Δ rrp6Δ* | BY4741 with *gcn5Δ::KanMX4 rrp6Δ::hph* | This work |
| LPY917 wild-type | *MATa ade2-101 his3Δ-200 leu2Δ1 trp1Δ1 lys2-801 TELadh4::URA3* | (6) |
| LPY917 *rrp6Δ* | LPY917 with *rrp6Δ::KanMX4* | This work |
| Nrd1-AA | *MATalpha tor1-1 fpr1::NAT RPL13A-2xFKB12::TRP1 Nrd1-FRB::kanMX6* | (7) |
| FSY1742 wild-type | *MATa ade2 his3 leu2 trp1 ura3* | (8) |
| FSY3117 *rrp6Δ* | FSY1742 with *rrp6Δ::KANr* | (8) |
| FSY3383 *rpd3Δ* | FSY1742 with *rpd3Δ::TRP1* | (7) |
| FSY3384 *rpd3Δ rrp6Δ* | FSY1742 with *rpd3Δ::TRP1 rrp6Δ::KANr* | (7) |
| Sth1-AA | *MATalpha tor1-1 fpr1::NAT RPL13A-2xFKB12::TRP1 Sth1-FRB::kanMX6* | (9) |
| Sth1-AA *gcn5Δ* | Sth1-AA with *gcn5Δ::SpHIS5* | This work |
| Sth1-AA *rrp6Δ* | Sth1-AA with *rrp6Δ::KanMX4* | This work |
| FSY6857 | *MATa bar1Δ::hisG BrdU-Inc::HIS3* | This work |
| FSY5439 | *MATα trp1::TRP1* | This work |
| FSY9286 | *MATa bar1Δ::hisG BrdU-Inc::HIS3 pho5Δ::URA3* | This work |

| | | |
|---|---|---|
| FSY9287 | *MAT$\alpha$ trp1::TRP1 pho5$\Delta$::URA3* | This work |
| FSY9288 | *MATa bar1$\Delta$::hisG BrdU-Inc::HIS3 PHO5-Sense-Terminator* | This work |
| FSY9291 | *MAT$\alpha$ trp1::TRP1 PHO5-Antisense-Terminator* | This work |

**Table S2. Primers**

| Primer ID | Sequence (5'→3') | Description |
|---|---|---|
| RRP6-Kan1 | GATAGACGAAATAGGAACAACAAACAGCTTATAAGCACCCAATAAGTGCGTTCCCGGCCAGCGACATGGAGGCCCAG | Deletion of *RRP6* (KanMX4 marker) |
| RRP6-Kan2 | GCCCTTGGTCCATTACTATCGCTAGATGATGGGTCGAATCTCCTTTTCCGAATCGACAGCAGTATAGCGACCAGGCT | Deletion of *RRP6* (KanMX4 marker) |
| RRP6hph_fwd | GAAATAGGAACAACAAACAGCTTATAAGCACCCAATAAGTGCGTTACGATCATTCAAGAGATCCCCG | Deletion of *RRP6* (hph marker) |
| RRP6hph_rev | AATTACCATAATTTATAAATAAAAAAATACGCTTGTTTTACATAATACCGCCTTTGAGTGAGCTG | Deletion of *RRP6* (hph marker) |
| RRP6_fwd | CCCAAAAATATGAGGGCATCGG | Confirmation of *RRP6* deletion |
| RRP6_rev | AAAATGGTGTGCATGGGGGA | Confirmation of *RRP6* deletion |
| RRP6ORF_fwd | AATAACCCCAGTCACTACCC | Confirmation of *RRP6* deletion |
| RRP6ORF_rev | ACAACACCGAAAACCTTTCC | Confirmation of *RRP6* deletion |
| gcn5HIS_fwd | GTGAGCCGCCCAAAAGTCTTCAGTTAACTCAGGTTCGTATTCTACATTAGCCGCTAGGGATAACAGGGT | Deletion of *GCN5* |
| gcn5HIS_rev | ATTTATTTCTTCTTCGAAAGGAATAGTAGCGGAAAAGCTTCTTCTACGCAAAGAGCGCCCAATACGCAAA | Deletion of *GCN5* |
| gcn5HIScheck1_fwd | TGGTAAGGGAAGACCGTGAG | Confirmation of *GCN5* deletion |
| gcn5HIScheck1_rev | TCGTCTCGCCGTACTAAACA | Confirmation of *GCN5* deletion |
| gcn5HIScheck2_fwd | GACAATGCCGCAAAAGTCCA | Confirmation of *GCN5* deletion |
| gcn5HIScheck2_rev | CCCGACAAGTCAACTACGCT | Confirmation of *GCN5* deletion |
| TEF1PHO5AS_fwd | AAAATTTGGGTATTCGTATTTAGTTTCCAATATTATTTAGTTATACAAAACGTACGCTGCAGGTCGAC | Insertion of *TEF1* promoter to drive *PHO5* AS transcription |
| TEF1PHO5AS_rev | ACTGGGACTGGAACACTACTCATTACAACGCCAGTCTATTGAGACAATAGCATCGATGAATTCTCTGTCG | Insertion of *TEF1* promoter to drive *PHO5* AS transcription |
| TEF1PHO5AS_check1_fwd | TGGGCAACACTTTCCACAGA | Confirmation of *TEF1-PHO5AS* construct |
| TEF1PHO5AS_check1_rev | TCCTGACTGACTACAGGGATTGA | Confirmation of *TEF1-PHO5AS* construct |

| | | |
|---|---|---|
| TEF1PHO5AS_check2_fwd | GGTGCCGGACCATACTACTC | Confirmation of *TEF1-PHO5AS* construct |
| TEF1PHO5AS_check2_rev | ACGGTCTTCAATTTCTCAAGTTTC | Confirmation of *TEF1-PHO5AS* construct |
| PMA1_fwd | CAATCTAATCACGGTGTCGACGACGAAGAC | RT-qPCR |
| PMA1_rev | GGCTTCCATAACGAATTGAATTGGACCG | RT-qPCR |
| SCR1_fwd | AACCGTCTTTCCTCCGTCGTAA | RT-qPCR |
| SCR1_rev | CTACCTTGCCGCACCAGACA | RT-qPCR |
| PHO5prom_qPCR_fwd | CACGTGGGACTAGCACAGAC | RT-qPCR |
| PHO5prom_qPCR_rev | TGCCTTGCCAAGTAAGGTGA | RT-qPCR |
| PHO5ORF_qPCR_fwd | ACACTCGTCAATTCAACGGC | RT-qPCR |
| PHO5ORF_qPCR_rev | GAGCATCCAGTGTATGGGTTCA | RT-qPCR |
| PHO5_5adjreg_fwd | CCTTTACCGTAATTTTCAATTGCTAA | ChIP qPCR |
| PHO5_5adjreg_rev | TCGCTTCTTCAACAGTGGTAAAAATA | ChIP qPCR |
| PHO5_UASp2_fwd | GAATAGGCAATCTCTAAATGAATCGA | ChIP qPCR |
| PHO5_UASp2_rev | GAAAACAGGGACCAGAATCATAAATT | ChIP qPCR |
| OFS2869 | TTTATAGGTTAAGGATAGTAAAGGAATACAGGTAAG | Plasmid construction |
| OFS2870 | ACTCACAAATTAGATAATTATCCTATAAATATAACGTTTTTGAACAC | Plasmid construction |
| OFS2871 | TAGGATAATTATCTAATTTGTGAGTTTAGTATACATGC | Plasmid construction |
| OFS2872 | TCCTTTACTATCCTTAACCTATAAAAATAGGCGTATCACGAG | Plasmid construction |
| OFS5084 | AGAACAACAACAAATAGAGCAAGCAAATTCGAGATTACCAcagctgaagcttcgtacgc | Strain construction |
| OFS5085 | TATTCGTATTTAGTTTCCAATATTATTTAGTTATACAAAAgcataggccactagtggatctg | Strain construction |
| OFS5086 | CATGAGAATAAGAACAACAACAAATAGAGCAAGCAAATTCGAGATTAGTAATGAATTAAGTCTTGATATATAACAATTAGCTTGATGTTTAAATCTGTTGTTTATTCAATTTTAGCCGC | Strain construction (Sense terminator Forward) |
| OFS5087 | CGGCAAAATTTAGATAAAAAATTTGGG | Strain construction (Sense terminator Reverse) |
| OFS5088 | GTTAGTATGGCTTCATCTCTCATGAGAATAAGA | Strain construction (Antisense terminator Forward) |
| OFS5089 | AAAAATTTGGGTATTCGTATTTAGTTTCCAATATTATTTAGTTATACAGTAATGAATTAAGTCTTGATATATAACAATTAGCTTGCTATTGTCTCAATAGACTGGCGTTGTAATG | Strain construction (Antisense terminator Reverse) |
| OFS2522_PHO5_fwd | CTTGGGACTACGATGCCAAT | RT-qPCR |
| OFS2523_PHO5_rev | ACTTCAAATGCACACCACGA | RT-qPCR |

| | | |
|---|---|---|
| OFS1717_SCR1_fwd | AACCGTCTTTCCTCCGTCGTAA | RT-qPCR |
| OFS1718_SCR1_rev | CTACCTTGCCGCACCAGACA | RT-qPCR |
